# A highly diverse set of novel immunoglobulin-like transcript (NILT) genes in zebrafish indicates a wide range of functions with complex relationships to mammalian receptors

**DOI:** 10.1101/2022.04.21.489081

**Authors:** Dustin J. Wcisel, Alex Dornburg, Sean C. McConnell, Kyle M. Hernandez, Jorge Andrade, Jill L. O. de Jong, Gary W. Litman, Jeffrey A. Yoder

**Author notes:** Kite Pharma, Santa Monica, CA 90404 USA.

## Abstract

Multiple novel immunoglobulin-like transcripts (NILTs) have been identified from salmon, trout and carp. NILTs typically encode activating or inhibitory transmembrane receptors with extracellular immunoglobulin (Ig) domains. Although predicted to provide immune recognition in ray-finned fish, we currently lack a definitive framework of NILT diversity, thereby limiting our predictions for their evolutionary origin and function. In order to better understand the diversity of NILTs and their possible roles in immune function, we identified five NILT loci in the Atlantic salmon (*Salmo salar)* genome, defined 86 NILT Ig domains within a 3 Mbp region of zebrafish (*Danio rerio*) chromosome 1, and described 41 NILT Ig domains as part of an alternative haplotype for this same genomic region. We then identified transcripts encoded by 43 different NILT genes which reflect an unprecedented diversity of Ig domain sequences and combinations for a family of non-recombining receptors within a single species. Zebrafish NILTs include a sole putative activating receptor but extensive inhibitory and secreted forms as well as membrane-bound forms with no known signaling motifs. These results reveal a higher level of genetic complexity, interindividual variation and sequence diversity for NILTs than previously described, suggesting that this gene family likely plays multiple roles in host immunity.

## Introduction

The ability of a multicellular organism, or host, to survive in pathogen-rich environments relies on the innate ability to differentiate between normal host cells (self) and pathogens or infected cells (non-self). A key component of a successful defense includes a wide range of cell surface immune receptors designed to recognize pathogens and abnormal host cells and target their destruction. Although certain innate immune receptors are highly conserved from vertebrates to invertebrates (e.g. TOLL-like receptors), vertebrates possess a number of gene families that encode clusters of rapidly evolving innate immune receptors that are not well conserved (e.g. natural killer receptor KIR and Ly49 families) (Yoder and Litman 2011). This rapid, lineage-specific diversification of innate immune receptors may be driven by historic pathogen exposures and host-pathogen co-evolution (Jack 2015; Papkou et al. 2019) that correspondingly has driven the evolution of lineage specific gene families. For example, work in teleost fishes has led to the description of multiple immunoglobulin (Ig) domain-containing innate immune receptor (IIIR) families including the novel immune-type receptor (NITR), diverse immunoglobulin-type receptor (DICP), leukocyte immune-type receptors (LITR), polymeric immunoglobulin receptor-like (PIGRL), and novel immunoglobulin-like transcript (NILT) families (Stet et al. 2005; Panagos et al. 2006; Stafford et al. 2006; Østergaard et al. 2010; Yoder and Litman 2011; Montgomery et al. 2011; Rodríguez-Nunez et al. 2014; Rodriguez-Nunez et al. 2016; Wcisel and Yoder 2016). A critical first step in understanding the evolutionary origins and molecular function of IIIR gene families is cataloging the diversity of these sequences within and between species.

Members of the IIIR family of NILTs have been reported in common carp (*Cyprinis carpio*), rainbow trout (*Oncorhynchus mykiss*) and Atlantic salmon (*Salmo salar*), which includes membrane bound proteins possessing one or two Ig ectodomains (Stet et al. 2005; Kock and Fischer 2008; Østergaard et al. 2009, 2010; Rodríguez-Nunez et al. 2014). The majority of all Ig domains, including those of NILTs, contain a pair of cysteines, C^23^ and C^104^ [numbering based on the IMGT system (Lefranc et al. 2003)] that play a role in stabilizing the Ig fold (Harpaz and Chothia 1994). All NILT Ig domains reported to date also possess an additional cysteine pair, internal to the canonical C^23^ and C^104^ pair, that are separated by 3, 6 or 7 residues (Cx_3_C, Cx_6_C, or Cx_7_C motifs) (Østergaard et al. 2010). As observed for many IIIR families (Rodríguez-Nunez et al. 2014; Wcisel and Yoder 2016), the majority of NILTs possess either cytoplasmic immunoreceptor tyrosine-based inhibitory motifs (ITIM: S/I/V/LxYxxI/V/L), ITIM-like sequences (itim: YxxI/V/L) or an immunoreceptor tyrosine-based activation motif (ITAM: YxxI/Lx_6-12_YxxI/L), and thus are predicted to provide inhibitory or activating immune function (Taylor et al. 2000; Barrow and Trowsdale 2006; Lanier 2008; Bezbradica and Medzhitov 2012). The expression of NILTs in carp and trout is highest in immune tissues (spleen, gill, thymus, kidney and gut) when compared to non-immune tissues (e.g. muscle, liver and heart) (Stet et al. 2005; Kock and Fischer 2008) and at least one NILT is upregulated in carp macrophages after infection (Joerink et al. 2006)further suggesting a role in immune function.

The most detailed genomic analysis of a NILT gene cluster was described from Atlantic salmon (prior to the completion of a reference genome) with the sequencing of a ∼200 kbp BAC that encodes six NILT genes and two NILT pseudogenes (Østergaard et al. 2010). Initial efforts to mine the zebrafish (*Danio rerio*) reference genome indicated that zebrafish encode multiple NILTs on chromosome 1 (Stet et al. 2005), and, although two zebrafish genomic sequences were named *nilt1* and *nilt2* (Østergaard et al. 2009), no other studies have been reported on the genomic organization of NILT gene clusters in any species. Given that NILT Ig domains have been shown to share sequence and structural homology with human Natural Cytotoxicity Triggering Receptor 2 (NCR2, aka NKp44) and Triggering Receptor Expressed on Myeloid Cells (TREM) receptors, as well as members of the CD300 (aka CMRF35) family (Stet et al. 2005; Østergaard et al. 2009), extensively documenting their diversity is critical to determine if they potentially represent orthologs to these mammalian IIIRs.

The completion of reference genomes for zebrafish (Howe et al. 2013) and Atlantic salmon (Lien et al. 2016) now makes detailed genomic analyses and syntenic comparisons of NILT loci possible. Here we define the chromosomal location of five different NILT loci in salmon and provide evidence for conserved synteny between salmon NILT loci with a single NILT gene cluster on zebrafish chromosome 1. In depth transcriptomic and genomic sequence comparisons of NILTs between individual zebrafish and between zebrafish lines reveal; predicted NILT proteins with 1, 2, 4 and 6 Ig domains, unprecedented sequence diversity between NILT Ig domains, and intraspecific gene content variation within this gene family.

Although the presence of ITIMs and ITAMs in NILTs is conserved from salmonids (salmon and trout) to cyprinids (zebrafish and carp), the sequence, number, and organization of Ig domains within individual receptors are not well conserved, indicating lineage-specific NILT gene cluster diversification. These observations indicate that the cellular signaling component of NILTs has been largely conserved for several hundred million years (Near et al. 2012; Dornburg et al. 2014; Dornburg and Near 2021), while the ligands recognized by NILTs have likely evolved recently within teleost lineages. These observations illustrate that NILTs are a highly diverse, “fish-specific” IIIR gene family, and our sequence data and analyses described herein reveal a heretofore unappreciated level of genomic and transcriptional complexity. Collectively, our results provide further evidence of unique pathways of taxa-specific variation within this family of immune-type receptors and the necessary framework for future studies investigating their ligands and functions.

## Methods

### Animals

All experiments involving live zebrafish were performed in accordance with relevant institutional and national guidelines and regulations, and were approved by the North Carolina State University Institutional Animal Care and Use Committee. Zebrafish strains included TU, AB, EKW, NHGRI-1, LSB, HSB, FS, CG1, and CG2. TU (Tübingen) zebrafish were a gift from John Rawls (Duke University). The AB line was procured through the Zebrafish International Resource Center (ZIRC). EKW zebrafish were purchased from EkkWill Waterlife Resources (Ruskin, FL). FS zebrafish were purchased from Drs. Foster & Smith (Rhinelander, Wisconsin). The NHGRI-1 line was derived from select individuals of the highly polymorphic TU line (Haffter et al. 1996; LaFave et al. 2014) and a gift from Shawn Burgess (National Human Genome Research Institute). LSB and HSB individuals were three generations derived from wild-caught zebrafish (Wong et al. 2012) and gifts from John Godwin (NC State University). CG1 and CG2 are genetically distinct, homozygous diploid clonal lines derived from *golden* zebrafish (Mizgirev and Revskoy 2014) and gifts from Sergei Revskoy (University of Kentucky College of Medicine).

### Nomenclature for NILTs

Throughout this manuscript, genus-species abbreviations are employed when needed to differentiate between NILTs from different species [e.g. zebrafish (*Danio rerio*) *nilt1* = *Dare-nilt1*; salmon (*Salmo salar*) *NILT1* = *Sasa-NILT1*]. However, when discussing only zebrafish NILTs, no genus-species abbreviations are provided.

Zebrafish *nilt1* (GenBank BN001234) and *nilt2* (GenBank BN001235) sequences were previously reported (Østergaard et al. 2009). Additional zebrafish NILT sequences identified in this study were named consecutively *nilt3, nilt4, nilt5* etc. Putative pseudogenes were assigned the next number in the symbol series and suffixed by a “p”. Sequences were considered pseudogenes if an exon encoding a NILT Ig domain possessed an internal stop codon or if no transcripts were identified for a genomic NILT Ig domain. This nomenclature system for zebrafish NILT sequences was designed in consultation with, and approved by, the Zebrafish Nomenclature Coordinator (https://zfin.org/).

### Acquisition and analyses and Atlantic salmon NILT sequences

Previously published NILT sequences were obtained from the literature (Stet et al. 2005; Kock and Fischer 2008; Østergaard et al. 2009, 2010) and are summarized in **Supplementary Table S1**. Additional Atlantic salmon NILT sequences were identified from predicted protein sequences from the Atlantic salmon reference genome (ICSAG2_v2; RefSeq GCF_000233371.1) via BLASTp searches (E-value <1e-8) using all previously reported NILT sequences as queries (Altschul et al. 1990) and included in **Supplementary Table S1**.

Protein structures of published and newly identified Atlantic salmon NILTs were identified using SMART software (Letunic and Bork 2018) and NILT identity confirmed by assessing the evolutionary relationships of Ig domains with established NILT Ig domains aligned by Clustal Omega v1.2.4 (Sievers and Higgins 2021). Evolutionary relationships were estimated using maximum likelihood in IQ-TREE 2 (Minh et al. 2020), with the amino acid substitution model identified as the best-fitting under Bayesian Information Criterion (Cavanaugh 2016) and 1000 ultrafast bootstrap approximation replicates to quantify clade support (Hoang et al. 2018). BoxShade (version 3.21) alignment plots were made using Clustal Omega output and manually annotated (https://embnet.vital-it.ch/software/BOX_form.html).

### Syntenic Analysis of Atlantic Salmon and Zebrafish Genomes

Protein sequences from the annotated Atlantic salmon genome (GCF_000233371.1) were used in reciprocal BLASTp searches against Ensembl GRCz11 zebrafish proteins (Release 101) with a maximum of five returned hits at an E-value of <1e-8. To ensure genes with multiple isoforms were only represented by one protein sequence, only the first protein sequence encountered in the annotation file for each genome was used in the BLAST analysis using custom scripts in Perl. The results of the BLAST searches between Atlantic salmon and zebrafish were subjected to collinear analyses using MCScanX (Wang et al. 2012). Collinear links (syntenic blocks) between chromosomes were extracted and formatted for visualization by modifying the Perl scripts used for Collinearity (v1.0; https://github.com/reubwn/collinearity).

Modified scripts and additional custom scripts utilized to extract and format data can be found at https://github.com/djwcisel/nilts. Collinear links between chromosomes, defined by matching a minimum number of five genes within the span of 25 genes with a conservative MCScanX score of ≤ 25 (default 50), were visualized using Circos v0.69-9 (Krzywinski et al. 2009).

### Sequence coverage of alternative zebrafish genomes

Zebrafish chromosome 1 from GRCz11 was modified by placing the 202,236 bp sequence of Zv9-scaffold100 between adjacent contigs GRCz11-CTG104 and GRCz11-CTG111. Genomic coverage of alternative zebrafish assemblies, CG1 (NCBI BioProject [PRJNA454110](https://www.ncbi.nlm.nih.gov/bioproject/454110)) and CG2 (NCBI BioProject [PRJNA454109](https://www.ncbi.nlm.nih.gov/bioproject/454109)) (McConnell et al. 2016), against this modified chromosome were visualized using MUMmer4 (Marçais et al. 2018). Genomic coverage of alternative genome assemblies was calculated by summing the alignment length from MUMmer results after removing overlapping segments and dividing by the total reference sequence length. Percent identities were estimated by averaging the percent identity of alignments with overlaps.

### Genomic identification of zebrafish NILT immunoglobulin domains

The zebrafish reference genomes [11th (GRCz11), 10th (GRCz10) and 9th (Zv9)] were searched for NILT Ig domains using BLAST and all previously reported NILTs sequences as queries. As non-rearranging Ig domains are typically encoded in single exons, a six-frame translation of relevant contigs and scaffolds was performed and all open reading frames longer than 65 amino acids subjected to HMMER v3.3 analysis against the SMART database (Eddy 2001; Letunic and Bork 2018). After the initial pass, a hmm profile was constructed using identified NILT Ig domains and the six-frame translation was searched again with HMMER. A reciprocal BLASTp search was used to exclude Ig domains that returned only hits to annotated zebrafish B-cell receptor CD22-like sequences in GRCz11. Predicted NILT Ig domains (E-value < 0.001) and any overlap with genes annotated by NCBI or Ensembl pipelines are summarized in **Supplementary Table S2**.

### Analyses of Ig Domains

Alignments of Ig domains were performed using the default settings of Clustal Omega and manually refined by eye to ensure proper alignment of the Cx_n_C motifs. Evolutionary relationships were inferred using identical methods to those described above. NILT Ig domains were grouped according to their internal cysteine motifs: Cx_3_C, Cx_6_C, Cx_7_C (Østergaard et al. 2010) or atypical if the Cx_n_C motif was absent and no clear group could be identified. For each of the four NILT groups, BoxShade alignment plots were generated following methods outlined above.

### Zebrafish Transcriptome Analyses

In order to assist the annotation of the genes encoding the NILT Ig domains, we mined existing transcriptome data from immune-related tissues (kidney, intestine, gills and spleen) of homozygous diploid clonal CG2 zebrafish (NCBI accession number SRP057116) (Dirscherl and Yoder 2015) and generated new RNAseq data from homozygous diploid clonal CG1 zebrafish. Tissues from six individual CG1 zebrafish were pooled and RNA was isolated using TRIzol (Invitrogen) following the manufacturer’s protocol. Quantity, purity and integrity of RNA was estimated using a NanoDrop 1000 (Thermo Fisher) and a TapeStation (Agilent). Equal amounts of RNA from each tissue type were pooled for a final concentration of 180 ng/μl and an RNA integrity number (RIN) of 7.3. Library preparation and next-gen sequencing was performed by Novogen Corporation Inc (Sacramento, CA) on a NovaSeq 6000 instrument (Illumina). Adapter sequences and poor quality reads were filtered with Trimmomatic v34 (Bolger et al. 2014). The transcriptome was *de novo* assembled with Trinity v2.11.0 (Grabherr et al. 2011). Raw reads and computationally assembled transcriptome sequences were deposited onto NCBI under the project accession number PRJNA672972.

Zebrafish CG1 and CG2 transcriptomes were searched using tBLASTn and all zebrafish genomic NILT Ig domains as queries (summarized in **Supplementary Table S2**). Putative NILT transcripts were then used as BLASTn queries to search the reference genome and retained if top hits aligned to the NILT regions of zebrafish chromosome 1, filtering out gene models belonging to the B-cell receptor CD22-like family that is also present in this genomic region. Transcripts were translated *in silico* and predicted protein domains were identified using SMART. Inhibitory signaling motifs (e.g., ITIM/itim) were identified by manual inspection of protein sequences.

### Visualization of genome and transcriptome sequences

Integrative Genomics Viewer (IGV 2.4.1) was used to visualize the genomic context of NILT Ig domains, Ensembl and NCBI annotations, alternative genomes and transcriptomes (Robinson et al. 2011). CG1 and CG2 genomic sequences (NCBI BioProjects [PRJNA454110](https://www.ncbi.nlm.nih.gov/bioproject/454110) and [PRJNA454109](https://www.ncbi.nlm.nih.gov/bioproject/454109)) (McConnell et al. 2016) were aligned to the modified chromosome 1 sequence using BWA v0.7.17 with a seed length of 2000 bp (Li and Durbin 2009). NILT transcripts identified from CG1 and CG2 transcriptomes were mapped using Splign v2.1.0 (Kapustin et al. 2008).

### Genotyping NILT Haplotypes

A PCR-based genotyping approach was used to confirm the presence of two NILT haplotypes, hap100 and hap111, in different zebrafish strains. PCR primers were designed to amplify a 313 bp fragment of genomic DNA unique to hap111 (111-Forward: TCCATATAATGCTAAATATGAGGA, 111-Reverse: CTCTGTGTTTATTGCAGTAATTGA).

Separate primers were designed to amplify a 186 bp fragment of hap100 (100-Forward: AGGAAAATGCTCAATAGGTTAT, 100-Reverse: GTCATCAAATTTTCCCAGTCTT). Genomic DNA was prepared using fin-clips from adult zebrafish or 1 day post-fertilization embryos following the hot sodium hydroxide method (Truett et al. 2000). A ‘touchdown’ PCR protocol was employed with GoTaq polymerase (Promega) consisting of an initial cycle of 95 °C for 30 seconds, 60 °C for 30 seconds, and 72 °C for 30 seconds. For the subsequent 19 cycles, the annealing temperature was decreased 0.5 °C each cycle. Finally, 30 cycles were executed with an annealing temperature of 50 °C. PCRs were carried out individually and amplicon sequences confirmed by Sanger sequencing. Haplotype amplicons were combined for agarose gel electrophoresis. As a positive control, sequences of the ribosomal first internal transcribed spacer region (ITS1) were PCR amplified using the same PCR conditions above with previously described primers [ITS1-F: AAAAAGCTTTTGTACACACCGCCCGTCGC, ITS1-R: AGCTTGCTGCGTTCTTCATCGA (Pleyte et al. 1992)].

## Results and Discussion

### Genome analysis of Atlantic salmon reveals multiple NILT genes and loci

Prior to this study a total of fourteen NILTs had been described from teleosts: two from common carp (*Cyca-NILT1, Cyca-NILT2*), four from rainbow trout (*Onmy-NILT1, Onmy-NILT2, Onmy-NILT3, Onmy-NILT4*), six from Atlantic salmon (*Sasa-NILT1, Sasa-NILT2, Sasa-NILT3, Sasa-NILT4, Sasa-NILT5, Sasa-NILT6*) and two predicted from zebrafish genomic sequence (*Dare-nilt1, Dare-nilt2*) (Stet et al. 2005; Kock and Fischer 2008; Østergaard et al. 2009, 2010). All previously reported NILT sequences contain one or two Ig ectodomains (termed D1 and D2 with D1 being membrane distal and D2 membrane proximal), are typically membrane bound, and possess cytoplasmic ITIMs or a cytoplasmic ITAM (**Fig. 1a**). It should be noted that due to recent lineage-specific diversification of the NILT family, one-to-one NILT orthologs are not identifiable between species, and NILT gene names reflect only the order in which they were identified. This is emphasized by the observation that, as a group, carp NILTs are only ∼30-60% identical to salmon NILTs (Stet et al. 2005; Østergaard et al. 2010), and this relationship can be observed in **Fig. 1b** where a phylogenetic analysis demonstrates that *Sasa-NILT1*, for example, is not a “true” ortholog of either *Onmy-NILT1* or *Cyca-NILT1*.

**Fig. 1.**
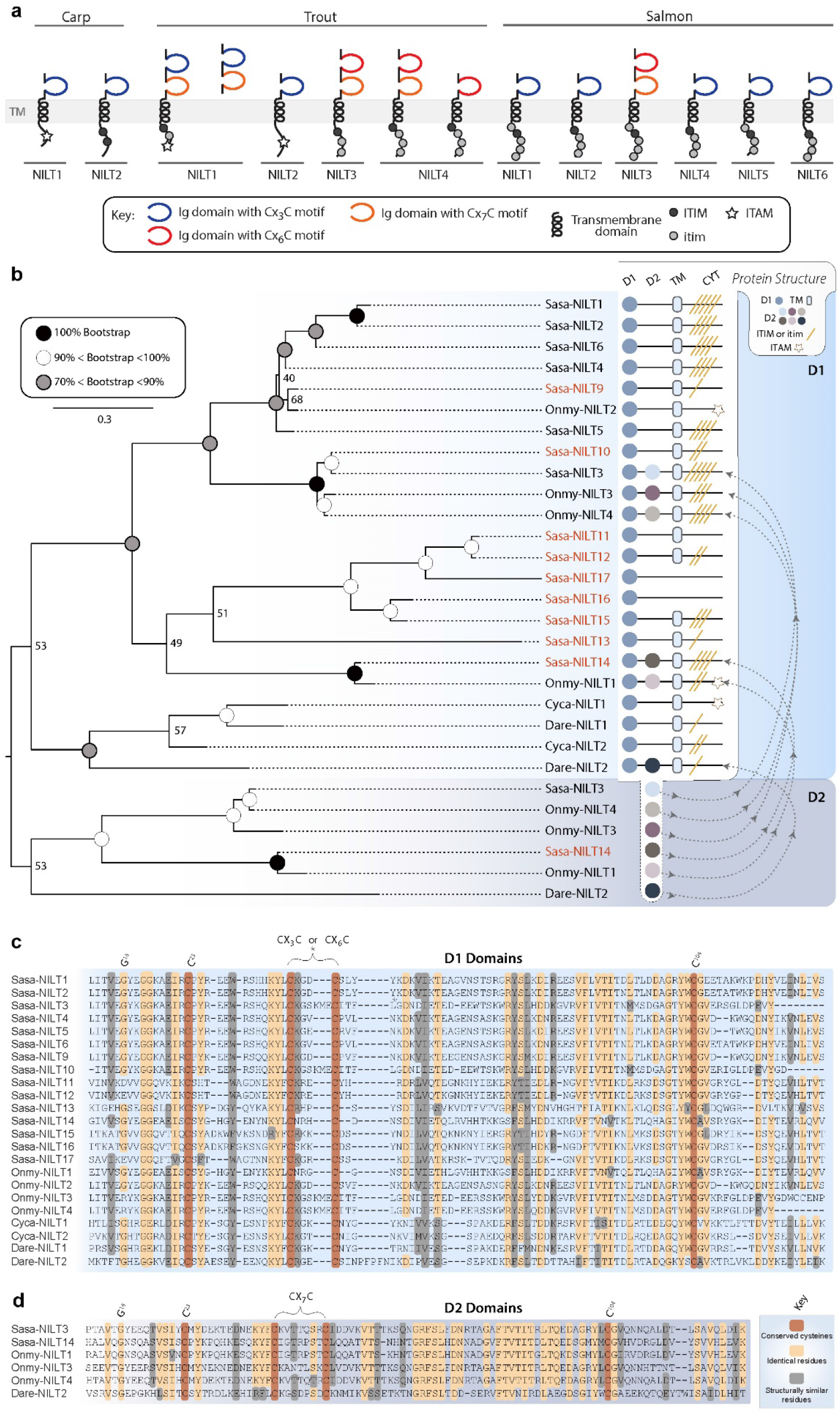
Sequence comparisons of previously reported and newly identified Atlantic salmon NILTs. **a** Protein architecture of previously reported NILTs (Stet et al. 2005; Kock and Fischer 2008; Østergaard et al. 2009, 2010). **b** Evolutionary relationships of NILT immunoglobulin (Ig) domains (D1 and D2) were estimated using maximum likelihood in IQ-TREE 2. Previously reported NILTs including the predicted Dare-Nilt1 and Dare-Nilt2 were compared to salmon NILT sequences identified in this study (brown text). Previously reported partial Sasa-NILT7 and Sasa-NILT8 sequences (Østergaard et al. 2010) were excluded from this analysis. Bootstrap support is indicated at each node. **a-b** Predicted protein structures include: D1 = 1st Ig domain, D2 = 2nd Ig domain, TM = transmembrane domain, CYT= Cytoplasmic tail, ITIM = immunoreceptor tyrosine-based inhibitory motif, itim = ITIM-like sequence, ITAM = immunoreceptor tyrosine-based activation motif. **c-d** Alignments of NILT D1 and D2 domains. Highly conserved key residues are numbered based on the IMGT system (Lefranc et al. 1999). Light orange shading indicates residues which are identical in >80% of sequences. Gray shading indicates similar residues conserved across sequences. Conserved cysteines are highlighted with dark orange. Dashes indicate gaps in the alignment. Species identifiers and gene names: Atlantic salmon (*Salmo salar; Sasa*), rainbow trout (*Oncorhynchus mykiss; Onmy*), common carp (*Cyprinus carpio L*.; *Cyca*), and zebrafish (*Danio rerio; Dare*). Sequences and identifiers (Genbank) are listed in **Supplementary Table S1**.

In the initial report describing six salmon NILT genes (*Sasa-NILT1* through *Sasa-NILT6*), three additional, but partial, NILT Ig domains were identified by genomic PCR and named *Sasa-NILT7, Sas-NILT8* and *Sasa-NILT9* (Østergaard et al. 2010). A search of the Atlantic salmon reference genome using previously published NILT Ig domains as queries identified *Sasa-NILT1* through *Sasa-NILT6*, a gene corresponding to the *Sasa-NILT9* amplicon, and eight additional NILT genes, *Sasa-NILT10* through *Sasa-NILT17* (**Fig. 1b and Supplementary Table S1**). *Sasa-NILT9, Sasa-NILT10, Sasa-NILT12, Sasa-NILT13, Sasa-NILT14*, and *Sasa-NILT15* possess cytoplasmic ITIM or itim sequences and are predicted to provide an inhibitory function.

Predicted sequences for *Sasa-NILT11, Sasa-NILT16* and *Sasa-NILT17* lack exons encoding a cytoplasmic tail prohibiting the prediction of their signaling potential. All salmon NILT D1 domains possess a Cx_3_C or Cx_6_C motif, whereas the D2 domains possess a Cx_7_C motif (**Fig. 1c-d**). A phylogenetic analysis of Ig domains from these new salmon NILTs along with published NILTs reveals distinct clusters for D1 and D2 domains. The D1 and D2 clades become further resolved when salmonid or cyprinid families are considered (**Fig. 1b**). This lineage-specific clustering is indicative of recent gene birth and death events followed by rapid diversification (Nei and Rooney 2005; Fernández and Gabaldón 2020) and is observed for other multigene families of Ig domain-containing innate immune receptors, e.g. NITRs (Desai et al. 2008).

### Atlantic salmon NILT11 shares conserved synteny with zebrafish nilt1 and nilt2

A high-quality genome assembly of Atlantic salmon has since become available after the initial description of six NILT genes from a BAC clone (Østergaard et al. 2010; Lien et al. 2016). The BAC containing these genomic NILT sequences, as well as *Sasa-NILT9* and *Sasa-NILT10*, can be readily identified on Atlantic salmon chromosome 3. Four additional NILTs, *Sasa-NILT11, Sasa-NILT12, Sasa-NILT13* and *Sasa-NILT14*, appear to exist as single copy genes on chromosomes 2, 6, 13, and 19, respectively. *Sasa-NILT15, Sasa-NILT16* and *Sasa-NILT17* are on unplaced genomic scaffolds. Our analyses revealed no large-scale conserved synteny exists between the cluster of Atlantic salmon NILTs on chromosome 3 and the single copy NILTs found on other chromosomes (**Fig. 2**). However, the region of chromosome 2 that encodes *Sasa-NILT11* appears to share a number of syntenic relationships with a region of zebrafish chromosome 1 that encodes *Dare-nilt1* and *Dare-nilt2*. While a large amount of synteny exists between the region of chromosome 6 that encodes *Sasa-NILT12* and chromosome 3, NILTs were not detected in this region of chromosome 3.

**Fig. 2.**
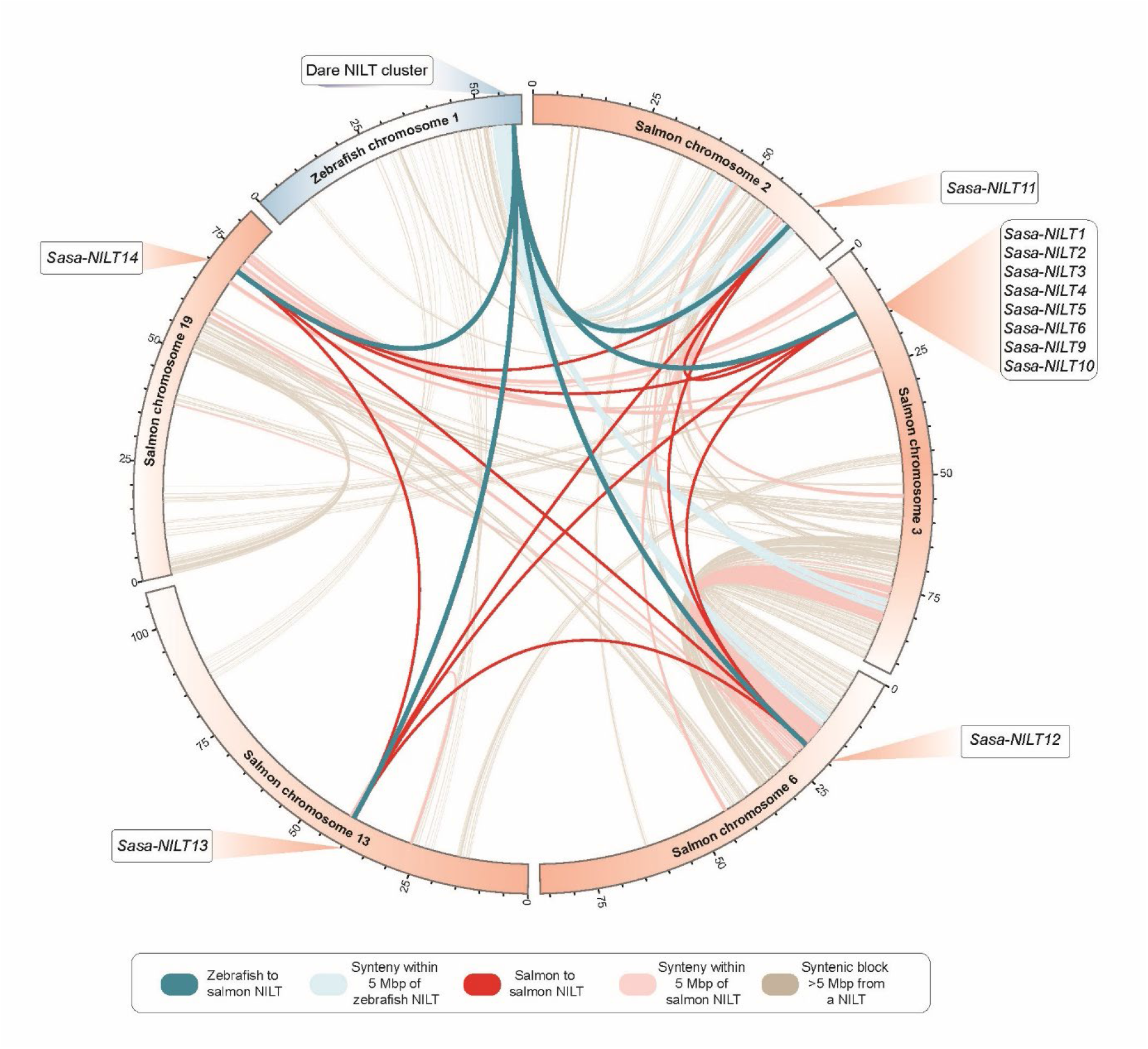
NILT loci in Atlantic salmon chromosomes 2, 3, 6, 13 and 19 and zebrafish chromosome 1 and their syntenic relationships. Approximate positions of NILT loci on a chromosome (salmon= orange; zebrafish = blue) are indicated by pop-outs. Dark blue lines connect zebrafish NILTS to salmon NILTs. Light blue lines connect syntenic blocks < 5 Mbp from a zebrafish NILT to salmon chromosomes 2, 3, 6, 13, and 19. Dark red lines connect salmon to salmon NILT locations. Light red lines connect syntenic blocks < 5 Mbp from a salmon NILT to salmon chromosomes 2, 3, 6, 13 and 19. Gray lines indicate syntenic blocks that are > 5 Mbp from a NILT.

### Diverse zebrafish NILT immunoglobulin domains are clustered on Chromosome 1

Initial BLAST searches of the 11th (GRCz11), 10th (GRCz10) and 9th (Zv9) builds of the zebrafish reference genome using published NILT sequences as queries revealed numerous hits to chromosome 1 on contigs GRCz11-CTG111 (NW_018394163.1), GRCz11-CTG104

(NW_018394162.1), GRCz10-CTG112 (NW_001878090.4), Zv9-scaffold101 (NW_001878090.3) and Zv9-scaffold100 (NW_003040340.2). Zv9-Scaffold100 was retired and removed from both GRCz10 and GRCz11. In contrast, Zv9-scaffold101 was incorporated into GRCz10-CTG112 which was then incorporated into GRCz11-CTG111; therefore, Zv9-scaffold100 (hereafter referred to as scaffold100) was the only non-GRCz11 sequence used in downstream analyses. We identified a number of NILT Ig domains in scaffold100 that did not map to CTG104, CTG111 or anywhere else in the current (GRCz11) reference genome. A sequence alignment of scaffold100 and CTG111 reveals numerous extended segments of similarity as well as a number of gaps in the alignment suggesting that these sequences reflect structural variation of the same genomic locus (**Supplementary Fig. S1** and see below).

All identified zebrafish NILT Ig domains were combined to generate a hmmer profile that was used to reanalyze the zebrafish reference genome GRCz11 and Zv9-scaffold100 for additional NILT Ig domains (**Fig. 3a**). This analysis identified 86 NILT Ig domains in the GRCz11 reference genome with all of them present on chromosome 1: 83 on CTG111 and 3 on CTG104. In addition, 41 NILT Ig domains were identified on scaffold100 (**Fig. 3b** and **Supplementary Table S2**). In order to annotate these sequences, neighboring Ig domains were considered to be part of the same gene if this was supported by transcripts (**Fig. 3c-f** and **Supplementary Table S3**, see below) and Ig domains within a gene were numbered based on their relative position within the predicted protein (the D1 Ig domain is closest to the amino terminus). Genomic Ig domains lacking an identifiable transcript or possessing a premature stop codon were named pseudogenes.

**Fig. 3.**
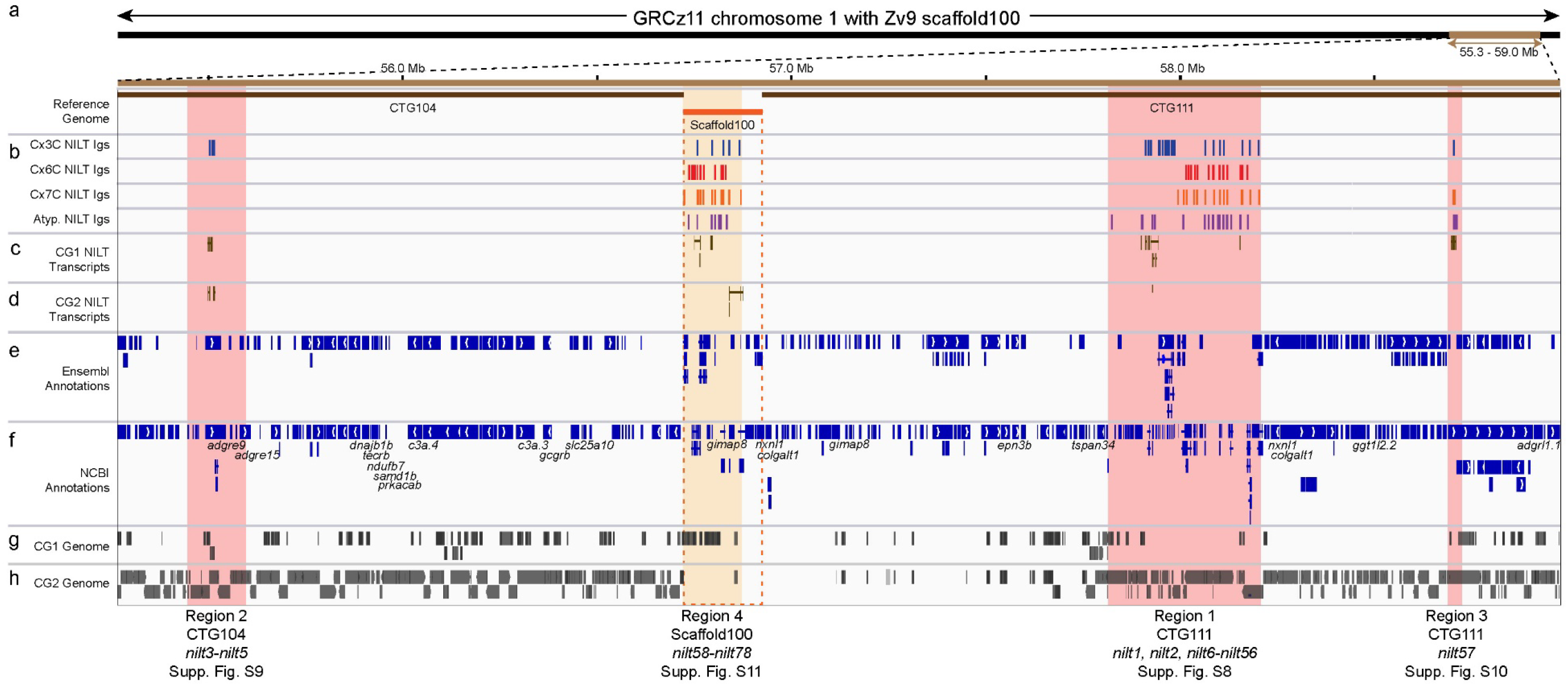
Genomic organization and variations within the zebrafish NILT gene cluster. Integrated Genomics Viewer display of the NILT gene cluster region of **a** GRCz11 chromosome 1 with Zv9 scaffold100 (202,236 bp) inserted between CTG104 and CTG111 (starting at 56,705,447 bp) encoding the NILT gene cluster. **b** Adjusted positions of NILT Ig domains, color-coded according to Cx_n_C groups. Splign alignments of NILT transcripts from **c** CG1 and **d** CG2 zebrafish. **e** Gene annotations from Ensembl v104 and scaffold 100 Ensembl v79. **f** RefSeq gene annotations from NCBI (NC_007112.7) and scaffold 100 (NW_003040340.2). *De novo* assembled genomic sequences from **g** CG1 and **h** CG2 zebrafish lines were mapped onto the modified GRCz11 chromosome reference genome. NILT regions 1 - 4 are defined at the bottom and displayed in more detail in **Supplementary Figs. S8 - S11**.

Phylogenetic analyses of these 127 Ig domains fail to define definitive groups of NILT Ig domains (**Supplementary Fig. S2**). This is in contrast to the clearly defined D1 and D2 domains of NILTs reported from other teleosts (**Fig. 1b**) (Østergaard et al. 2010). Indeed, a phylogenetic comparison of all zebrafish NILT Ig domains with those from salmon, trout and carp demonstrates that the sequence diversity of NILTs is far greater in zebrafish than expected (**Supplementary Fig. S3**). Our search of the Atlantic salmon reference genome identified only 19 NILT Ig domains and 17 NILT genes (**Supplementary Table S1**). For comparison, genomic PCR from a single carp revealed 53 different NILT Ig domains (Stet et al. 2005) suggesting that cyprinids (carp and zebrafish) may have experienced a larger lineage-specific expansion and diversification of NILTs than salmonids.

NILT Ig domains previously have been categorized based on the presence of a Cx_3_C, Cx_6_C or Cx_7_C motif which accounted for all reported NILTs from salmon, trout and carp (Stet et al. 2005; Kock and Fischer 2008; Østergaard et al. 2009, 2010). In contrast, only ∼70% (88 of 127) of zebrafish NILT Ig domains possess a Cx_3_C, Cx_6_C or Cx_7_C motif (**Supplementary Fig. S4-S6**). The remaining 39 Ig domains in this genomic region considered to be NILT Ig domains are named “atypical” (**Supplementary Fig. S7**). Interestingly, Ig domains within these groups do not always share the highest sequence similarities -- for example, the phylogenetic comparison of all zebrafish NILT Ig domains reveal that Cx_6_C and Cx_7_C domains are not reciprocally monophyletic (**Supplementary Fig. S2**). Instead Cx_6_C and Cx_7_C domains repeatedly arose independently, thereby raising the possibility that the difference between a 6 and a 7 amino acid inter-cysteine spacer does not impose a strong constraint for sequence conservation or that currently unknown selective pressures have driven patterns of convergent evolution.

### Zebrafish genome and transcriptome sequences reveal a highly expanded family of NILT genes

The genomic coordinates of the 127 zebrafish NILT Ig domains were used to search ENSEMBL and NCBI gene annotations in order to identify NILT genes. Seventy-one of the Ig domains are associated with 33 predicted genes in Ensembl and NCBI (**Supplementary Table S2**). Transcriptome sequence from immune tissues of CG1 and CG2 zebrafish, confirmed many of these predicted genes and identified ten additional NILT genes that were not annotated in Ensembl or NCBI (**Supplementary Table S3**). Overall 65 NILT transcripts were identified from CG1 and CG2 with some transcripts representing different alleles, RNA splice variants or, in some cases, truncated transcripts (**Supplementary Table S3**). In total, 43 NILT genes were identified in CTG104, CTG111, and scaffold100.

We identified three distinct NILT regions within the GRCz11 reference genome and one NILT gene cluster that is not present in the current reference genome but likely reflects a structural variant. Within GRCz11, there are: one major NILT gene cluster that is encoded in CTG111, spans ∼390 kbp and encodes 53 NILT genes/pseudogenes (“Region 1” in **Fig. 3**), one minor NILT gene cluster in CTG104 that encodes 3 NILT genes (“Region 2” in **Fig. 3**), and a minor NILT locus that encodes a single NILT gene (“Region 3” in **Fig. 3**). The major NILT gene cluster in Zv9-scaffold100 (“Region 4” in **Fig. 3**) spans ∼150 kbp, encodes 21 NILT genes/pseudogenes and is predicted to represent a structural variant of Region 1.

### Zebrafish NILT gene cluster includes four regions reflecting structural variants of chromosome 1

The major NILT gene cluster in GRCz11, Region 1, includes the previously described *nilt1* and *nilt2* genes (Østergaard et al. 2009) as well as 26 other NILT genes and 25 pseudogenes (**Fig. 3** and **Supplementary Fig. S8**). Although transcripts corresponding to *nilt1* and *nilt2* have not been reported previously, we identified *nilt2* transcripts from both NCBI and the CG2 transcriptome. The number of pseudogenes in this region is not unexpected for what is likely a recently expanding gene cluster - for example, twelve atypical Ig domains spanning ∼50 kb appear to be tandemly duplicated with other NILT Ig domains, but have no corresponding transcripts: *nilt31p, nilt32p, nilt33p, nilt34p, nilt35p, nilt36p, nilt38p, nilt39p, nilt41p, nilt42p, nilt47p*, and *nilt48p* (**Supplementary Fig. S8**). It is possible that some sequences annotated as pseudogenes may be transcribed under certain conditions (e.g. during infection) which can be reclassified as genes as needed.

The NILT Region 2 includes a total of three Ig domains that represent three NILT genes (*nilt3, nilt4* and nilt5). Although current ENSEMBL and NCBI annotations do not predict *nilt3* and *nilt4*, they are present in the CG1 and/or CG2 transcriptomes (**Supplementary Table S3**). We identified a single NCBI sequence corresponding to the *nilt5* Ig domain that is defined as a noncoding RNA (Genbank XR_002459137); in contrast, our CG2 *nilt5* transcript is predicted to encode a single Ig domain NILT with a transmembrane domain and cytoplasmic ITIM and itim sequences (**Supplementary Table S3**). One unusual feature of this minor NILT cluster is that *nilt3, nilt4* and *nilt5* are all encoded within a single intron of the *adgre9* gene (**Supplementary Fig. S9)**.

The NILT Region 3 includes six Ig domains that represent a single NILT gene (*nilt57*). Although ENSEMBL and NCBI do not predict any transcript that includes these Ig domains, we identified a *nilt57* transcript from CG2 that encodes all six Ig domains with a transmembrane domain and cytoplasmic ITIM and itim sequences (**Supplementary Fig. S10**).

The NILT Region 4 is encoded on Zv9-scaffold100 and predicted to reflect a structural variant of Region 1 in GRCz11-CTG111. Thirty of the 41 Ig domains in this region correspond to 21 NILT genes that are supported by NCBI, ENSEMBL and/or transcriptome sequences as well as 11 pseudogenes (**Supplementary Fig. S11**). Further support for Region 4 being a structural variant of Region 1 includes the presence of *nxnl, colgalt1* and *gimap8* genes in both scaffold100 and CTG111 (**Fig. 3f**). This structural variation between Region 1 and Region 4 is emphasized with differences in NILT gene content, reflective of haplotypic variation.

### CG1-specific NILT transcripts suggest an alternative gene content haplotype

Two NILT transcripts from CG1 zebrafish (bottom of **Supplementary Table S3**) did not map to the GRCz11 reference genome nor scaffold100 raising the possibility of gene content variation between haplotypes of this region of chromosome 1. Thus, we compared the whole genome assemblies of both CG1 and CG2 to the reference genome GRCz11. At the level of whole chromosomes, CG1 and CG2 do not appear to differ significantly in their genome coverage (**Fig. 4** and **Table 1**), suggesting accurate sequencing and assembly of both alternate genome assemblies. However, closer inspection of three specific, NILT-encoding regions within chromosome 1 of the reference genome (GRCz11) as well as scaffold100 (Zv9), reveal significant coverage differences between the CG1 and CG2 genome assemblies. CG1 genomic coverage to these NILT regions reveals a general lack of common sequences (**Table 1** and **Fig. 3g-h**). Although CG2 coverage is high for two NILT regions from GRCz11, low coverage is observed for the NILT region on scaffold100, suggesting CG2 encodes a NILT locus more similar to the GRCz11 structural variant, and that scaffold100 is absent from the genome of CG2. CG1 genomic coverage is much lower across three of the four regions, suggesting dramatic structural variations between CG1 and the reference genome exists. Further, the CG1 genome likely encodes a novel NILT gene-content haplotype. Indeed, this region of chromosome 1 has been shown to display an exceptional level of copy number variation (Brown et al. 2012; Rodríguez-Nunez et al. 2014).This identification of CG1 NILT transcripts which are not present in the reference genome is further evidence for structural and gene-content variation for the NILT gene cluster and is comparable to the variation observed for other zebrafish immune gene clusters such as the MHC Class I, NITR, and DICP gene clusters (Rodríguez-Nunez et al. 2014; Dirscherl and Yoder 2015; Rodriguez-Nunez et al. 2016; McConnell et al. 2016; Honjo et al. 2021).

**Table 1.** Coverage and percent identity of alternative zebrafish genome assemblies compared to GRCz11 reference genome.

| **Genomic Content** | **Contig/**<br>**Scaffold** | **Genomic**<br>**Coordinates<sup>i</sup>** | **Region in**<br>**Figure 4** | **CG1**<br>**Genomic**<br>**Coverage** | **CG1**<br>**percent**<br>**Identity** | **CG2**<br>**Genomic**<br>**Coverage** | **CG2**<br>**percent**<br>**Identity** |
| --- | --- | --- | --- | --- | --- | --- | --- |
| 25 Chromosomes | - | - | - | 87.23% | 96.43% | 87.91% | 96.50% |
| GRCz11 chromosome 1<br>with Zv9 scaffold100<sup>i</sup> | - | - | - | 88.48% | 96.58% | 89.46% | 96.74% |
| *nilt1, nilt2, nilt6 – nilt56* | CTG111 | 57797367-58189758 | Region 1 | 47.26% | 94.03% | 89.11% | 99.81% |
| *nilt3, nilt4, nilt5* | CTG104 | 55427288-55577739 | Region 2 | 67.45% | 95.24% | 93.81% | 99.96% |
| *nilt57* | CTG111 | 58670034-58707646 | Region 3 | 94.86% | 97.49% | 25.38 | 99.98% |
| *nilt58 – nilt78* | scaffold100 | 56703879-56854330 | Region 4 | 73.19% | 96.03% | 5.03% | 99.20% |

<sup>i</sup> Modified zebrafish chromosome 1 sequence presented in Figure 3.

**Fig. 4.**
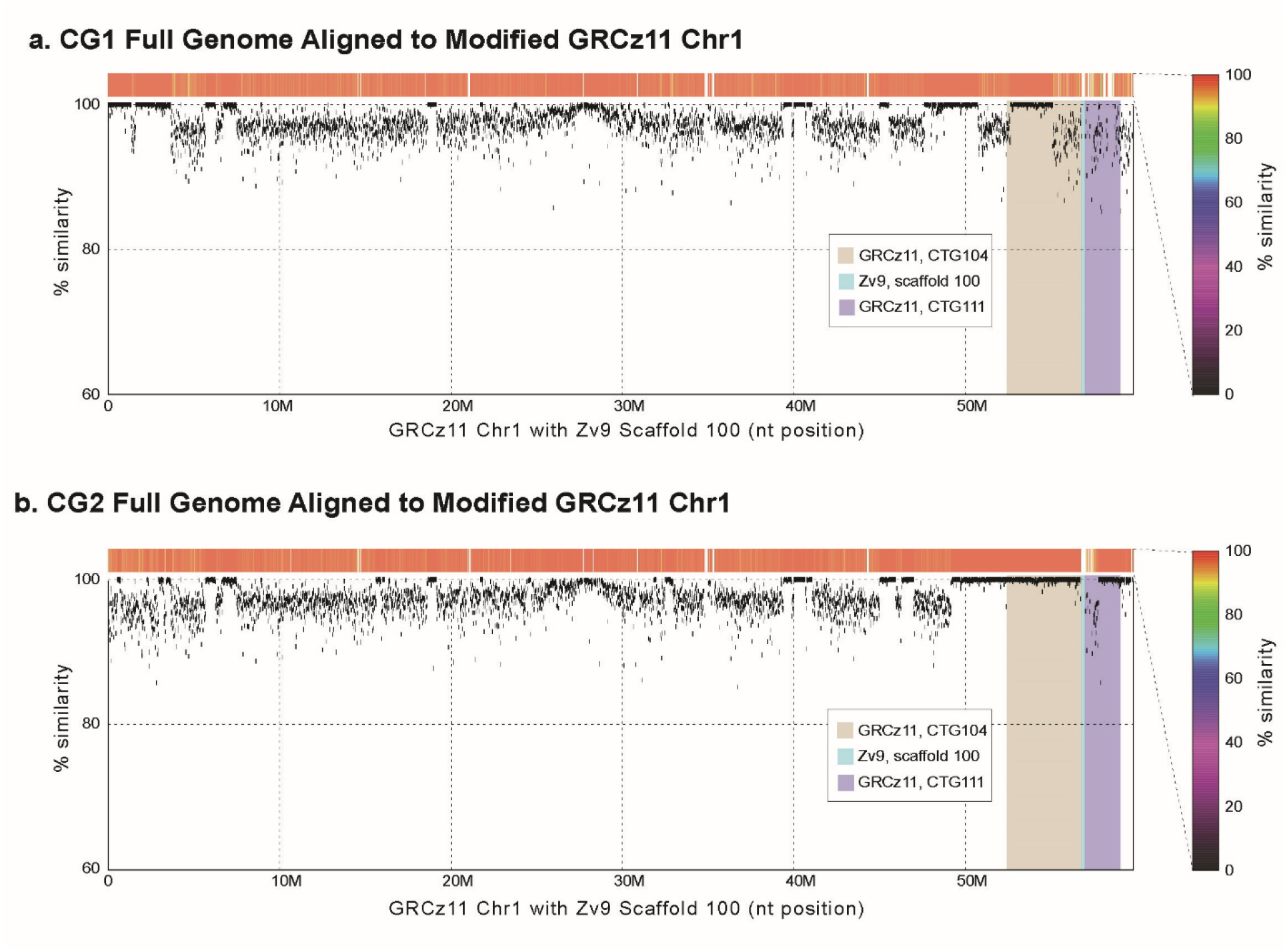
Coverage plots of alternative genome assemblies aligned to zebrafish chromosome 1. Scaffold 100 of Zv9 was inserted between CTG104 and CTG111 of GRCz11 beginning at base pair 56,705,447. **a** Scaffolds of de novo assembled CG1 genome mapped to modified zebrafish chromosome 1. **b** Scaffolds of de novo assembled CG2 genome mapped to modified zebrafish chromosome 1. Percent identity is shown as a heat map (above) and dotplot (below).

### A genotyping strategy confirms alternative gene content variation for the major NILT region in zebrafish

In order to demonstrate that Region 1 and Region 4 reflect different haplotypes for the same genomic region, a genotyping strategy was utilized with individuals from a range of zebrafish lines. PCR primers were designed to amplify sequences unique to Region 1 on CTG111 (hereafter referred to as hap111) and Region 4 on scaffold100 (hereafter referred to as hap100). Homozygous (hap100/100 or hap111/111) and heterozygous (hap100/111) individuals were identified from 9 different lines of zebrafish (**Fig. 5a**). CG1 was the only line of zebrafish where this genotyping strategy could not resolve either haplotype. The low coverage of the CG1 genome to these regions (**Table 1**) and two novel CG1 transcripts that do not map to the reference genome (**Supplementary Table S2**) indicates that CG1 reflects an additional structural variant of this genomic region that displays further gene content variation.

**Fig. 5.**
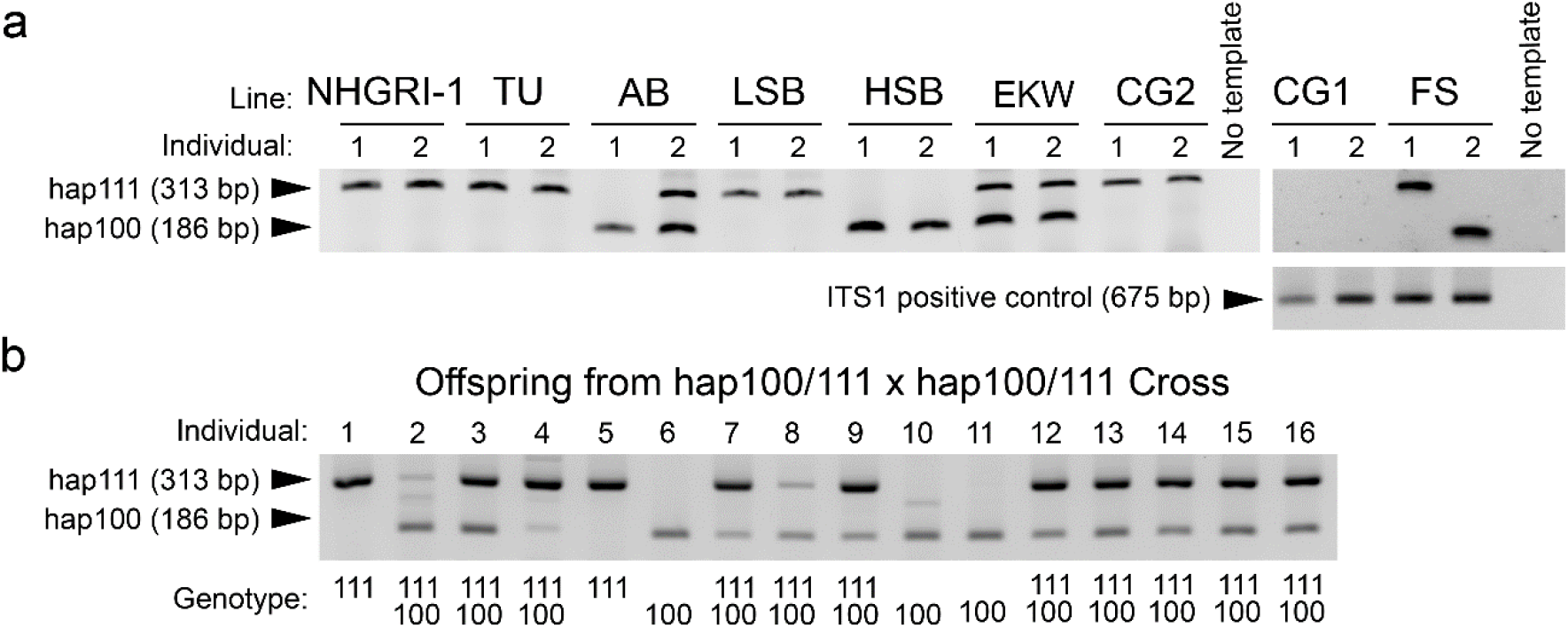
Genotyping confirms two NILT haplotypes in zebrafish. (**a**) Two individuals from nine wild-type zebrafish lines (NHGRI-1, TU, AB, LSB, HSB, EKW, FS, CG1 and CG2) were genotyped for the presence of NILT haplotypes. Genomic DNA (gDNA) from each individual was PCR amplified for hap100 and hap111, combined, and visualized by gel electrophoresis. ITS amplicons are shown as a positive control for CG1 which was negative for hap100/hap111 genotyping. (**b**) Sixteen offspring from two heterozygous (hap100/111) EKW zebrafish were genotyped revealing three homozygous hap100/100, eleven heterozygous hap100/111 offspring, and two homozygous hap111/111 individuals. This approximates the expected ratio of 1:2:1 for simple Mendelian inheritance of a single locus (see **Supplementary Fig. S12**).

Heterozygous (hap100/111) male and female zebrafish identified from a population of EKW zebrafish were crossed. The genotypes of the offspring were in the approximate ratio of 1:2:1 (hap100/100:hap100/111:hap101/111). These frequencies are consistent with frequencies expected by Mendelian inheritance of a single locus (Fig. 5b and Supplementary Fig. S12), and suggest that Region 4 (hap100 from Zv9) represents an alternative haplotype of Region 1 (hap111 from GRCz11). Given this observation, evidence that CG1 zebrafish encode a third NILT haplotype, and that this region of chromosome 1 displays significant copy number variation, we predict that multiple alternative haplotypes exist for the NILT gene cluster. Thus, there is a critical need for additional future work to assess the level of this variation, how such gene content variation arises, and the functional implications of this variation.

### Zebrafish NILT proteins display a larger degree of sequence variation than salmon, trout and carp

An examination of the predicted protein sequences of all zebrafish NILT genes and transcripts reveals a wide range of numbers and combinations of Ig domains (**Fig. 6** and **Supplementary Fig. S13**). Zebrafish NILTs possess 1, 2, 4 or 6 Ig domains and there are no obvious “rules” for the order or combinations of these domains within a protein. Single-Ig domain NILTs may possess a Cx_3_C, Cx_6_C, Cx_7_C or atypical Ig domain. All four types of Ig domains can be found at the amino terminus of a NILT. Importantly, atypical Ig domains are present in NILTs that also possess Cx_3_C, Cx_6_C, or Cx_7_C domains validating their inclusion in the repertoire of NILT Ig domains. One observed feature for multi-Ig domain NILTs is that Cx_3_C domains are not found to be membrane proximal - although this may not be consistent in other currently undescribed haplotypes. This degree of variation in the number and combination of Ig domains within NILTs has not been described in other species. In fact, our searches of the Atlantic salmon genome only reveal NILTs that possess one or two Ig domains and all can be classified as the “D1” or “D2” domains previously reported in carp, trout and salmon. This diversification of the extracellular NILT structures may be novel to zebrafish and perhaps closely related species (e.g. carp), but only analyses of additional reference genomes can resolve this issue. The large degree of variation in Ig domains likely reflects a diverse range of possible ligands. Although it is expected that annotations of NILTs in additional species will be facilitated by these zebrafish sequences, targeted searches and manual annotations will likely be necessary to identify all NILTs in other teleosts.

**Fig. 6.**
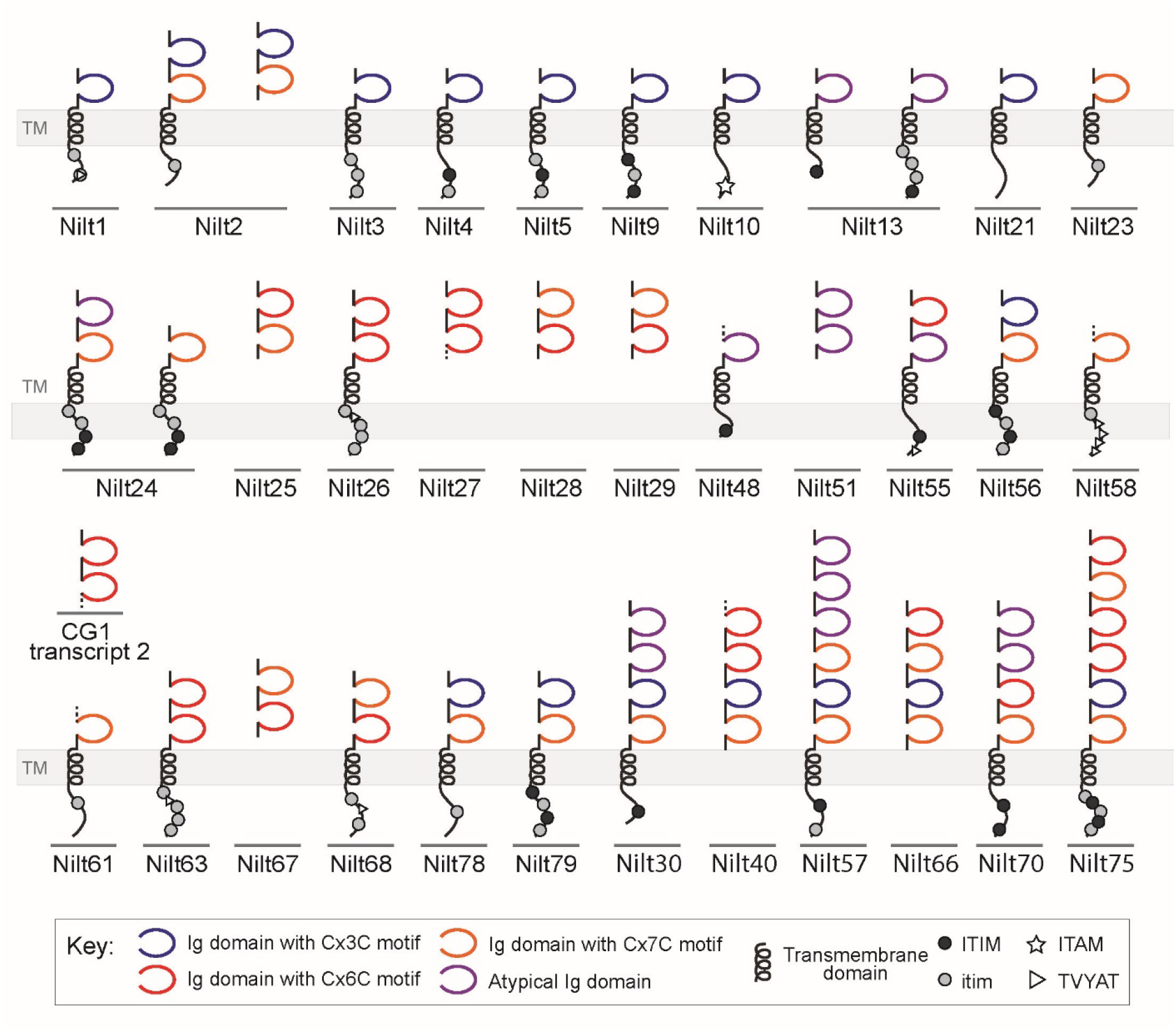
Highly diverse structures of zebrafish NILT proteins. Predicted protein structures of representative NILTs are shown (see **Supplementary Table S3** and **Supplementary Fig. S13**). Immunoglobulin domains are indicated by extracellular loops which are color coded based on the presence of a Cx_3_C (blue), Cx_6_C (red), or Cx_7_C (orange) motif or as an atypical (purple) sequence. Membrane bound NILTs include a transmembrane (TM) domain (e.g. Nilt1). NILTs lacking a TM domain are predicted to be secretory (e.g. Nilt25). Cytoplasmic immunoreceptor tyrosine-based inhibition motif (ITIM), ITIM-like (itim), immunoreceptor tyrosine-based activation motif (ITAM), and TVYAT sequences are represented by dark gray circles, light gray circles, white stars and white triangles, respectively.

The current collection of predicted zebrafish NILT sequences (**Fig. 6** and **Supplementary Fig. S13**) includes a single activating receptor (Nilt10) that possesses a classical cytoplasmic ITAM (**Supplementary Figs. S13** and **S14**), 25 inhibitory receptors that possess one or more cytoplasmic ITIM/itim (**Supplementary Figs. S13** and **S15**), 10 secreted proteins, and a single membrane bound NILT (Nilt21) with no obvious signaling motif. This pattern of receptors is comparable to that of other zebrafish innate immune receptor families, such as NITRs and DICPs, in which the number of inhibitory and ambiguous receptors significantly exceeds the number of activating receptors (Rodríguez-Nunez et al. 2014).

Our manual inspection of cytoplasmic domains for signaling motifs revealed a novel TVYAT peptide motif nine times in six different zebrafish NILTs (Nilt1, Nilt26, Nilt55, Nilt58, Nilt63 and Nilt68) and twice in Atlantic salmon (NILT3 and NILT5) (**Fig. 6** and **Supplementary Figs. S13** and **S16**). To our knowledge, TVYAT has no identified signaling potential, but it’s potential for tyrosine phosphorylation and conservation across species boundaries may suggest a functional role.

### On the origins of NILTs

Based on sequence similarities, protein modeling, and limited evidence for conserved synteny, the original description of NILTs suggested that they might be related either to *NCR2*(NKp44) and the TREM family of IIIRs on human chromosome 6 or to the CD300 family of IIIRs on chromosome 17 (Stet et al. 2005). BLAST searches at that time identified CD300C (CMRF35), CD300LF (IREM1), TREM1, TREM2 and NCR2 (NKp44) as the human proteins having the highest sequence homology to the carp NILTs. Current BLASTp searches of human proteins with these carp sequences identify CD300LF, which possesses a Cx_7_C motif, as the top hit for both carp NILT Ig domains (∼35% identity, ∼60% sequence homology and E-values of 2.00E-12 and 4.00E-15) (**Supplementary Table S4**). Because a wide range of sequence diversity is observed in our updated catalog of NILT Ig domains, we selected an additional 19 NILT Ig domains representing phylogenetically diverse sequences (**Supplementary Fig. S3**) to search the human protein database. These NILT Ig domains were used as queries for BLASTp searches of the NCBI human protein database. Our results reveal that different Ig domains had differing similarities to CD300s, NCR2 (NKp44), TREMs, Polymeric Immunoglobulin Receptor (PIGR) and Fc Fragment of IgA and IgM Receptor (FCAMR) as well as the Ig variable (V) domains of TCR and Ig genes (**Supplementary Table S4**). As NILTs are not known to undergo somatic recombination, and zebrafish TCR and Ig genes have been well defined on chromosomes 2, 3, 17 and 24 in zebrafish (Traver and Yoder 2020) we exclude a homology relationship between NILTs and TCR/Ig genes. Likewise, a two-Ig domain *pigr* gene encoding Cx_7_C motifs has been identified on zebrafish chromosome 2 (Kortum et al. 2014) and described in other teleosts [e.g. (Hamuro et al. 2007; Tadiso et al. 2011; Sheng et al. 2018)] indicating that, although they share this sequence motif, NILTs likely do not share a common origin with PIGR. This leaves the CD300 and NCR2(NKp44)/TREM gene clusters as the human IIIR families with the strongest relationship to NILTs based on sequence homology. However, it should be noted that the highest sequence identity between a NILT Ig domain and a human sequence, Sasa-NILT9-D1 and CD300LG, is only 42% with an E-value of 8.00E-21. In addition, BLASTp searches (using default settings) with certain NILT Ig domains failed to identify homology to any human protein (e.g. Dare-Nilt30-D1 and Dare-Nilt57-D2) (**Supplementary Table S4**).

Although earlier reports on NILTs emphasized the commonality of the Cx_n_C motifs between NILTS and NCR2(Nkp44) which possesses a Cx_7_C motif, it should be noted that nearly all human TREM and CD300 proteins along with PIGR also possess a Cx_6_C or Cx_7_C motif, although some possess a Cx_8_C (including FCAMR) or Cx_9_C motif (e.g. CD300LG). These observations make it clear that the evolutionary signal needed to strongly support the claim that the teleost NILTs are homologous to any one human protein family has eroded.

To further investigate possible links between NILTs and human IIIR families, we searched for evidence of conserved synteny between the zebrafish NILT gene cluster and the human genome. In the original description of NILTs (Stet et al. 2005), it was observed that NILT sequences were approximately 7 Mbp from a MHC class I “Z lineage” gene cluster on zebrafish chromosome 1 (Kruiswijk et al. 2002; Dirscherl and Yoder 2014) which is similar to the relationship of the TREM/NCR2(NKp44) gene cluster to the HLA on human chromosome 6. A reexamination of this linkage in zebrafish reveals that (in GRCz11) the zebrafish *mhc1zba, mhc1zca, mhc1zda* and *mhc1zea* gene cluster is encoded at Chromosome 1: 47,116,121 - 47,161,996, which is 8.3 Mbp from the closest NILT (*Dare-nilt3* at position 55,486,075 - 55,485,765).

With our detailed genomic analysis of the zebrafish NILT gene cluster we performed a more fine-detailed search for evidence of conserved synteny with the human genome and specifically, with human IIIR gene clusters. We find the strongest evidence of conserved synteny between the zebrafish NILT regions and human genome is between *dnajb1b, tecrb, ndufb7, samd1b, prkacab* (which are adjacent to the NILT Region 2) and *nxnl1* and *colgalt1* [which are within NILT Region 1 (hap111) *and* the alternative gene-content haplotype NILT Region 4 (hap100)] (**Fig. 3** and **Supplementary Figs. S8, S9** and **S11**), whose orthologs - *DNAJB1, TECR, NDUFB7, SAMD1, PRKACA, NXNL1* and *COLGALT1* - are encoded in human chromosome 19p13.11 - 19p13.12. This region of human chromosome 19p13 also encodes multiple Adhesion G Protein-Coupled Receptor E genes (*ADGRE2, ADGRE3, ADGRE5*) as observed at NILT Region 2 on zebrafish chromosome 1 (*adgre6, adgre7, adgre8, agre9*, etc).

Further, tandem duplicates of zebrafish complement 3a (*c3a*.*1, c3a*.*2, c3a*.*3*, etc) are present between NILT Regions 1 and 2 and the single copy human ortholog *C3* is encoded (next to *ADGRE1*) on 19p13.3. No major IIIR gene cluster is identifiable in 19p13 including the CD300 (chromosome 17), NCR2(NKp44)/TREM (chromosome 6) and *PIGR*/*FCAMR* (chromosome 1) gene families (**Supplementary Table S4**). It is noted that the leukocyte receptor complex (LRC) that encodes multiple IIIR families (e.g. KIRs, LILRs) is encoded on the other arm of human chromosome 19 (19q13), however, we find limited sequence homology between NILT Ig domains and those of the KIR and LILR families. These observations add to the complications of assigning homology or orthology of NILTs to specific human IIIR gene families and support the model that NILTs are too diverged from mammalian genes to assign homologs/orthologs.

### Conclusions

Here we define the complete zebrafish NILT repertoire encoded within a single 3.7 Mbp region of chromosome 1 from the GRCz11 reference genome that includes 30 genes and 25 pseudogenes. We also define part of an alternative haplotype for this gene cluster that encodes an additional 21 NILT genes and 11 pseudogenes. The intraspecific gene content variation of the NILT gene family may contribute to inter-individual differences in immune health complicating immune interventions in aquaculture species as the evolutionary advantage may lie in complexity rather than uniformity. This intraspecific genetic diversity may also confound immunological studies employing zebrafish as a model. Such differences are representative of those observed within the human population. Gene-content haplotypes, reminiscent of those identified here, have been associated with disease susceptibilities. For example, the human KIR gene family exhibits gene-content haplotypes (Misra et al. 2018) that may influence HIV progression (Martin and Carrington 2013) and mothers with certain combinations of KIRs (and their MHC ligands) are predisposed to preeclampsia (Hiby et al. 2004). Therefore, the contribution of NILT gene-content haplotypes to population health, as well as individual health, is of significant interest.

The number and variation of zebrafish NILT Ig domain sequences and NILT protein architecture is far more complex than the NILT sequences identified from the Atlantic salmon reference genome. This suggests that the NILT gene content may have experienced a lineage-specific expansion in cyprinids (e.g. zebrafish and carp) when compared to salmonids (salmon and trout). Further analyses of additional teleost whole genomes can resolve this possibility and also reveal if other lineage-specific expansions exist.

Since the initial description of two carp NILTs as being similar to human NKp44, TREM and CD300(CMRF35) receptors (Stet et al. 2005), NILTs have been repeatedly considered orthologues (Klesney-Tait et al. 2006; Clark et al. 2009; Flajnik et al. 2012; Gasiorowski et al. 2013), however, we did not uncover any evidence to substantiate this claim. On the contrary, our analysis suggests that NILTs are a recently and rapidly evolving gene cluster that, based on current information, is too far diverged from the common mammalian-teleost ancestor to make claims of homology or orthology between NILTs and human IIIRs. It will be of great interest to determine if authentic NILT orthologs can be identified in earlier diverging lineages such as holostei (gar and bowfin).

While ligands for multigene families of IIIRs are difficult to identify, future functional studies will reveal important features of NILTs. It should be noted, however, that the large number and sequence diversity of NILT receptors and the unknown nature of their ligands, will make designing such experiments challenging. Nevertheless, the presence of NILTs in cyprinids and salmonids indicates that they have been present for over 200 million years (Near et al. 2012; Dornburg et al. 2014) and likely play an important role(s) in immunity. As zebrafish are highly amenable to gene disruption (e.g. CRISPR) technologies, we predict that if NILTs play an important role in the immune response, reverse genetics studies partnered with immune challenges will quickly reveal a phenotype and lead to additional studies (Hwang et al. 2013; LaFave et al. 2014). The results described here further lay the foundation for functional and comparative studies which seek to explore the origins and evolution of vertebrate innate immunity.

## Acknowledgements

We thank Dereje Jima (North Carolina State University) for early discussions about zebrafish NILTs, James (Thomas) Howard (North Carolina State University) for technical support, and Shawn Burgess (National Human Genome Research Institute), John Godwin (North Carolina State University), David Langenau (Mass General Research Institute), John Rawls (Duke University) and Sergei Revskoy (University of Kentucky College of Medicine) for zebrafish lines.

## Funding

This work was supported by the National Science Foundation (IOS1755330 to JAY and IOS1755242 to AD), the National Institutes of Health (R01 AI057559 to GWL and JAY and R01 AI23337 to GWL), the National Evolutionary Synthesis Center, NSF EF0905606 (DJW), the Triangle Center for Evolutionary Medicine (AD and JAY), the Chicago Biomedical Consortium (CCT Searle Fund to Jd and SM), and by services provided by the University of Chicago Genomics Core Facility and Bioinformatics Core Facility which are supported by the UChicago Medicine Comprehensive Cancer Center NCI Cancer Support Center Grant (P30 CA014599).

## Data availability

New transcriptome sequence read data used in this study have been deposited to NCBI under the project accession number PRJNA672972. Custom Perl and R scripts used in the processing and formatting of data can be found on https://github.com/djwcisel/nilts

## Authors’ contributions

DW, GW and JY conceived the project; DW assembled transcriptome, datamined public databases and wrote custom scripts to facilitate these processes; AD performed phylogenetic analyses; SM, KH, JA and JdJ completed genome sequencing and assembly; DW, AD and JY collected and analyzed the data, created graphics and wrote the manuscript; JY supervised the project. All authors read and approved the manuscript.

## Conflict of interest

The authors declare that they have no conflicts of interest.

## Notes

### Competing Interest Statement

The authors have declared no competing interest.

