## Supplementary Figures for "A highly diverse set of novel immunoglobulin-like transcript (NILT) genes in zebrafish indicates a wide range of functions with complex relationships to mammalian receptors"

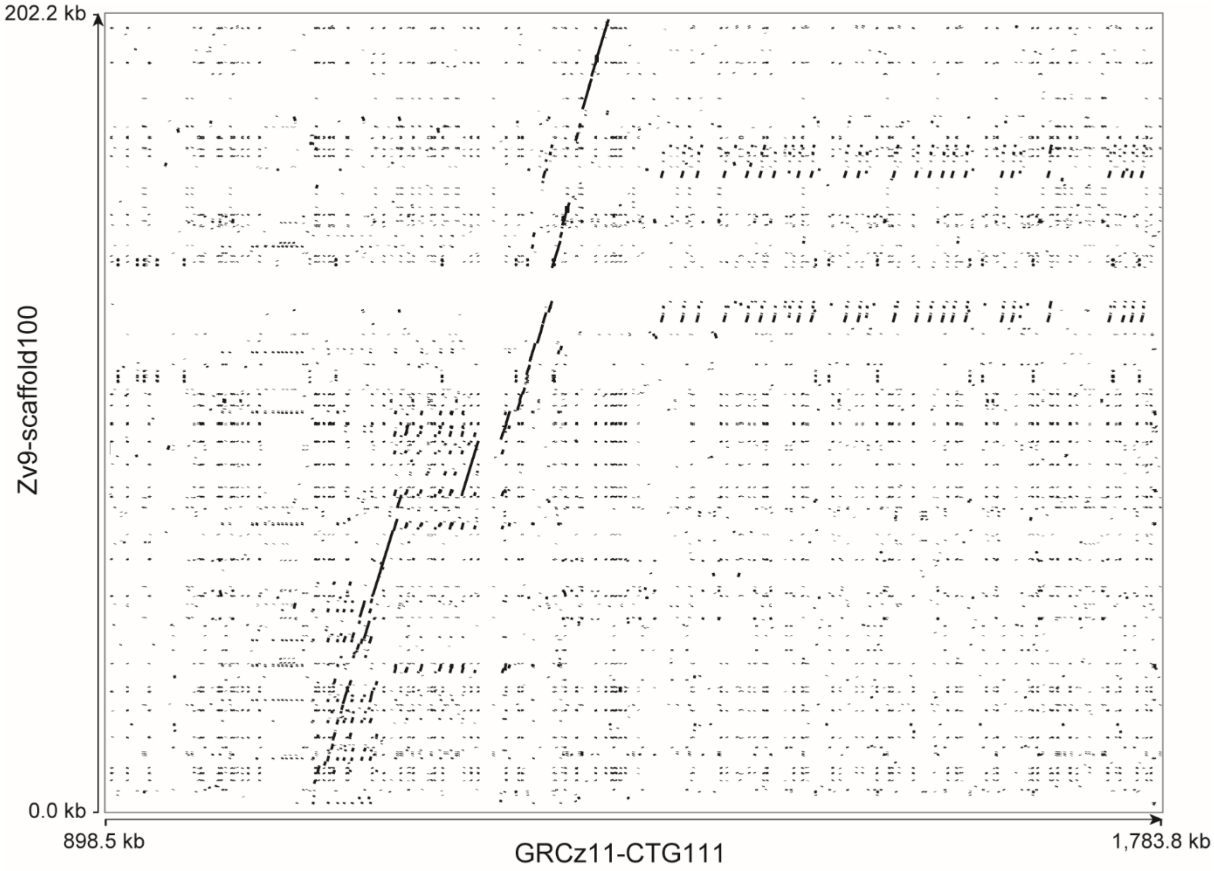

**Figure S1. Dot-plot sequence comparison of Zv9-scafffold100 to GRCz11-CTG111.**
The entire 202,232 bp sequence of Zv9-scaffold100 (NW_003040340.2) was aligned to 885,349 bp of GRCz11-CTG111 genomic coordinates (898,464-1,783,813 bp) corresponding to the start of the first NILT Ig domain (*nilt6p_D1*) and end of the last NILT Ig domain (*nilt57_D1*) using default settings of PipMaker (Schwartz et al. 2000).

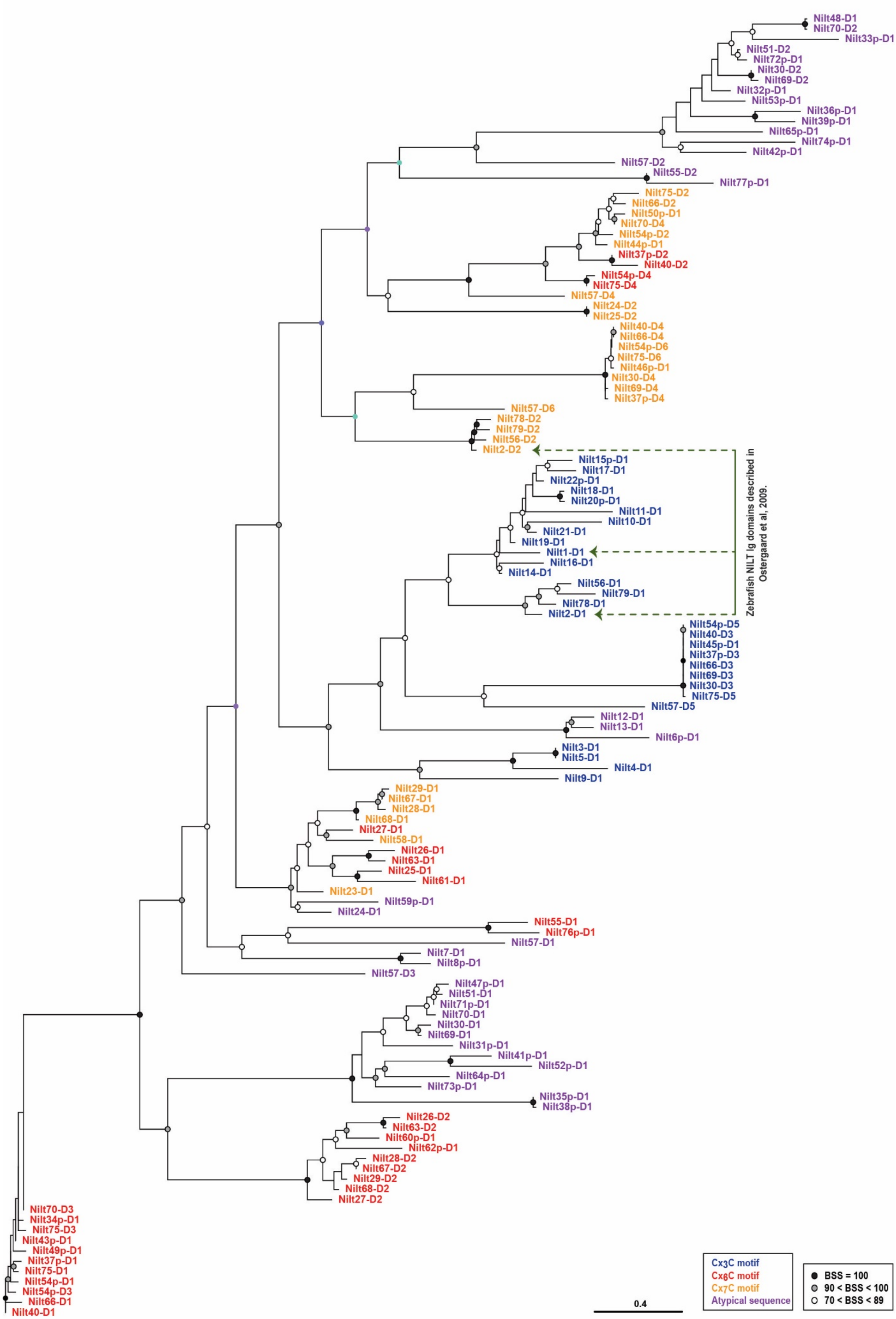
Nilt48-D1
Nilt70-D2
Nilt33p-D1
Nilt51-D2
Nilt72p-D1
Nilt30-D2
Nilt69-D2
Nilt32p-D1
Nilt53p-D1
Nilt36p-D1
Nilt39p-D1
Nilt65p-D1
Nilt74p-D1
Nilt42p-D1
Nilt57-D2
Nilt55-D2
Nilt77p-D1
Nilt75-D2
Nilt66-D2
Nilt50p-D1
Nilt70-D4
Nilt54p-D2
Nilt44p-D1
Nilt37p-D2
Nilt40-D2
Nilt54p-D4
Nilt75-D4
Nilt57-D4
Nilt24-D2
Nilt25-D2
Nilt40-D4
Nilt66-D4
Nilt54p-D6
Nilt75-D6
Nilt46p-D1
Nilt30-D4
Nilt69-D4
Nilt37p-D4
Nilt57-D6
Nilt78-D2
Nilt79-D2
Nilt56-D2
Nilt2-D2
Nilt15p-D1
Nilt17-D1
Nilt22p-D1
Nilt18-D1
Nilt20p-D1
Nilt11-D1
Nilt10-D1
Nilt21-D1
Nilt19-D1
Nilt1-D1
Nilt16-D1
Nilt14-D1
Nilt56-D1
Nilt79-D1
Nilt78-D1
Nilt2-D1
Zebrafish NILT Ig domains described in Ostergaard et al, 2009.
Nilt54p-D5
Nilt40-D3
Nilt45p-D1
Nilt37p-D3
Nilt66-D3
Nilt69-D3
Nilt30-D3
Nilt75-D5
Nilt57-D5
Nilt12-D1
Nilt13-D1
Nilt6p-D1
Nilt3-D1
Nilt5-D1
Nilt4-D1
Nilt9-D1
Nilt29-D1
Nilt67-D1
Nilt28-D1
Nilt68-D1
Nilt27-D1
Nilt58-D1
Nilt26-D1
Nilt63-D1
Nilt25-D1
Nilt61-D1
Nilt23-D1
Nilt59p-D1
Nilt24-D1
Nilt55-D1
Nilt76p-D1
Nilt57-D1
Nilt7-D1
Nilt8p-D1
Nilt57-D3
Nilt47p-D1
Nilt51-D1
Nilt71p-D1
Nilt70-D1
Nilt30-D1
Nilt69-D1
Nilt31p-D1
Nilt41p-D1
Nilt52p-D1
Nilt64p-D1
Nilt73p-D1
Nilt35p-D1
Nilt38p-D1
Nilt26-D2
Nilt63-D2
Nilt60p-D1
Nilt62p-D1
Nilt28-D2
Nilt67-D2
Nilt29-D2
Nilt68-D2
Nilt27-D2
Nilt70-D3
Nilt34p-D1
Nilt75-D3
Nilt43p-D1
Nilt49p-D1
Nilt37p-D1
Nilt75-D1
Nilt54p-D1
Nilt54p-D3
Nilt66-D1
Nilt40-D1
0.4
Cx3C motif
Cx6C motif
Cx7C motif
Atypical sequence
BSS = 100
90 < BSS < 100
70 < BSS < 89

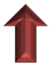

**Figure S2. Phylogenetic relationships among Ig domains of zebrafish NILTs.**
Genomic zebrafish NILT Ig domains were aligned by Clustal Omega (Sievers and Higgins, 2014). The resulting NILT domain alignment was analyzed using IQ-TREE 2 (Minh et al. 2020) to determine the best-fit model of amino acid substitution and then infer the evolutionary history of these domains using maximum likelihood. We used 1000 ultrafast bootstrap replicates (Hoang et al. 2018) to assess node support. Sequence names are color-coded based on the presence of Cx$_3$C (blue), Cx$_6$C (red) and Cx$_7$C (orange) motifs. Sequence names of atypical Ig domains are in purple text. The previously reported Ig domains present in zebrafish Nilt1 and Nilt2 (Østergaard et al., 2009) are indicated by arrows. BSS = % bootstrap support. Scale bar depicts branch lengths in substitution units.

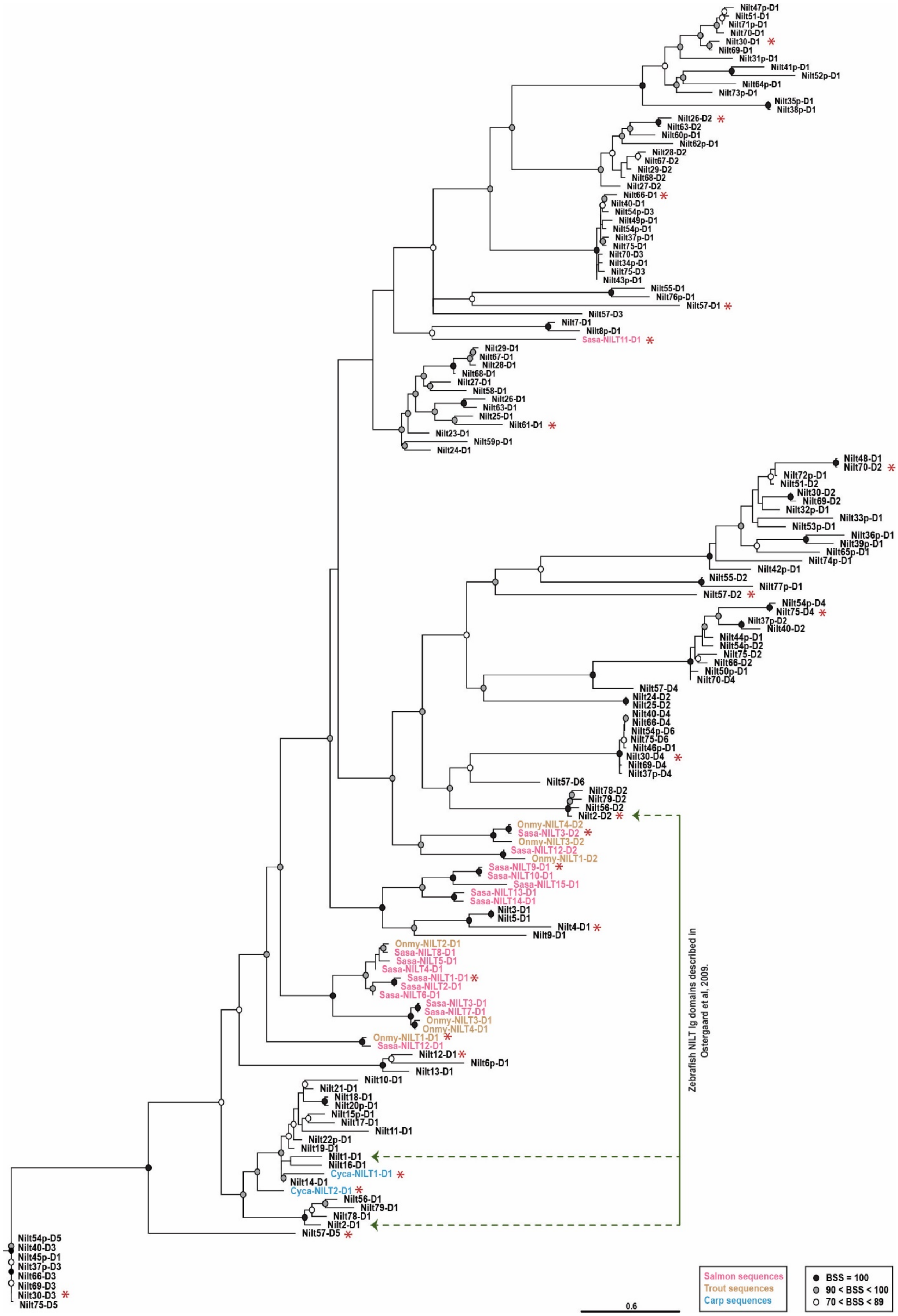

Nilt47p-D1
Nilt51-D1
Nilt71p-D1
Nilt70-D1
Nilt30-D1 *
Nilt69-D1
Nilt31p-D1
Nilt41p-D1
Nilt52p-D1
Nilt64p-D1
Nilt73p-D1
Nilt35p-D1
Nilt38p-D1
Nilt26-D2 *
Nilt63-D2
Nilt60p-D1
Nilt62p-D1
Nilt28-D2
Nilt67-D2
Nilt29-D2
Nilt68-D2
Nilt27-D2
Nilt66-D1 *
Nilt40-D1
Nilt54p-D3
Nilt49p-D1
Nilt54p-D1
Nilt37p-D1
Nilt75-D1
Nilt70-D3
Nilt34p-D1
Nilt75-D3
Nilt43p-D1
Nilt55-D1
Nilt76p-D1
Nilt57-D1 *
Nilt57-D3
Nilt7-D1
Nilt8p-D1
Sasa-NILT11-D1 *
Nilt29-D1
Nilt67-D1
Nilt28-D1
Nilt68-D1
Nilt27-D1
Nilt58-D1
Nilt26-D1
Nilt63-D1
Nilt25-D1
Nilt61-D1 *
Nilt23-D1
Nilt59p-D1
Nilt24-D1
Nilt48-D1
Nilt70-D2 *
Nilt72p-D1
Nilt51-D2
Nilt30-D2
Nilt69-D2
Nilt32p-D1
Nilt33p-D1
Nilt53p-D1
Nilt36p-D1
Nilt39p-D1
Nilt65p-D1
Nilt74p-D1
Nilt42p-D1
Nilt55-D2
Nilt77p-D1
Nilt57-D2 *
Nilt54p-D4
Nilt75-D4 *
Nilt37p-D2
Nilt40-D2
Nilt44p-D1
Nilt54p-D2
Nilt75-D2
Nilt66-D2
Nilt50p-D1
Nilt70-D4
Nilt57-D4
Nilt24-D2
Nilt25-D2
Nilt40-D4
Nilt66-D4
Nilt54p-D6
Nilt75-D6
Nilt46p-D1
Nilt30-D4 *
Nilt69-D4
Nilt37p-D4
Nilt57-D6
Nilt78-D2
Nilt79-D2
Nilt56-D2
Nilt2-D2 *
Onmy-NILT4-D2
Sasa-NILT3-D2 *
Onmy-NILT3-D2
Sasa-NILT12-D2
Onmy-NILT1-D2
Sasa-NILT9-D1 *
Sasa-NILT10-D1
Sasa-NILT15-D1
Sasa-NILT13-D1
Sasa-NILT14-D1
Nilt3-D1
Nilt5-D1
Nilt4-D1 *
Nilt9-D1
Onmy-NILT2-D1
Sasa-NILT8-D1
Sasa-NILT5-D1
Sasa-NILT4-D1
Sasa-NILT1-D1 *
Sasa-NILT2-D1
Sasa-NILT6-D1
Sasa-NILT3-D1
Sasa-NILT7-D1
Onmy-NILT3-D1
Onmy-NILT4-D1
Onmy-NILT1-D1 *
Sasa-NILT12-D1
Nilt12-D1 *
Nilt6p-D1
Nilt13-D1
Nilt10-D1
Nilt21-D1
Nilt18-D1
Nilt20p-D1
Nilt15p-D1
Nilt17-D1
Nilt11-D1
Nilt22p-D1
Nilt19-D1
Nilt1-D1
Nilt16-D1
Cyca-NILT1-D1 *
Nilt14-D1
Cyca-NILT2-D1 *
Nilt56-D1
Nilt79-D1
Nilt78-D1
Nilt2-D1
Nilt57-D5 *
Nilt54p-D5
Nilt40-D3
Nilt45p-D1
Nilt37p-D3
Nilt66-D3
Nilt69-D3
Nilt30-D3 *
Nilt75-D5
Zebrafish NILT Ig domains described in Ostergaard et al, 2009.
0.6
Salmon sequences
Trout sequences
Carp sequences
BSS = 100
90 < BSS < 100
70 < BSS < 89

**Figure S3. Phylogenetic relationships among Ig domains of zebrafish, salmon, trout and carp NILTs.**
Genomic zebrafish NILT Ig domains were aligned with previously reported Atlantic salmon, rainbow trout and common carp NILT Ig domains by Clustal Omega (Sievers and Higgins, 2014). The resulting NILT domain alignment was analyzed using IQ-TREE 2 (Minh et al. 2020) to determine the best-fit model of amino acid substitution and then infer the evolutionary history of these domains using maximum likelihood. We used 1000 ultrafast bootstrap replicates (Hoang et al. 2018) to assess node support. Sequence names are color-coded based on species: zebrafish (black), salmon (pink), trout (tan) and carp (light blue). The previously reported Ig domains present in Zebrafish Nilt1 and Nilt2 (Østergaard et al., 2009) are indicated by arrows. BSS = % bootstrap support. Scale bar depicts branch lengths in substitution units. Red asterisks indicate sequences used to search (BLASTp) the human protein database (see main text).

```
Nilt75-D5     VVRREGESAQIFCPYNSI-YQSKSKSLCKGKCSTR------DTRPLDETV-REEKERLTLHEDVTASVFTGTITGLTAEDAGKYWCAVTLDRELNYLYT
Nilt54p-D5    VVRREGESAQIFCPYDSI-YQSKSKSLCKGKCSTR------DTRPLDETV-REEKERLTLHEDVTASVFTGTITGLTAEDAGKYWCAVTLDRELNYLYT
Nilt30-D3     VVRREGESAQIFCPYDSI-YQSKSKSLCKGKCSTR------DTRPLDETV-REEKERLTLHEDVTASVFTGTITGLTAEDAGKYWCAVTLDRELNYLY-
Nilt37p-D3    VVRREGESAQIFCPYDSI-YQSKSKSLCKGKCSTR------DTRPLDETV-REEKERLTLHEDVTASVFTGTITGLTAEDAGKYWCAVTLDRELNYLYT
Nilt40-D3     VVRREGESAQIFCPYDSI-YQSKSKSLCKGKCSTR------DTRPLDETV-REEKERLTLHEDVTASVFTGTITGLTAEDAGKYWCAVTLDRELNYLYT
Nilt45p-D1    VVRREGESAQIFCPYDSI-YQSKSKSLCKGKCSTR------DTRPLDETV-REEKERLTLHEDVTASVFTGTITGLTAEDAGKYWCAVTLDRELNYLYT
Nilt66-D3     VVRREGESAQIFCPYDSI-YQSKSKSLCKGKCSTR------DTRPLDETV-REEKERLTLHEDVTASVFTGTITGLTAEDAGKYWCAVTLDRELNYLYT
Nilt69-D3     VVRREGESAQIFCPYDSI-YQSKSKSLCKGKCSTR------DTRPLDETV-REEKERLTLHEDVTASVFTGTITGLTAEDAGKYWCAVTLDRELNYLY-
Nilt57-D5     VMGREGDSAEIRCPNEDH-NLSEIKYFYKGKCFTQ-----EVPIIRSDEG-HIKRPHISMKDNTELNVFTVNISSLRAEDAGIYCCVVR----------
Nilt2-D1      -TGHEGGKVEIQCSYAESEYEENQKYLCRGECSINPFPFNIKDIPVESGS-EPKDKRFSLTDDTTAHIFTITITDLRTADQGKYSCAVKTRLV------
Nilt78-D1     -TGHEGGKVEIQCPYSESAYEGNQKYLCRGECPIKPFLFQNIDIPVESGS-APVDKRFSLTDNRTAHIFTITITDLRTADQGKYWCAVKTG--------
Nilt56-D1     -TGHEGGKVEIRCPYN-AKYEEMEKYLCRGECPIY-----DKDIPVESGS-EPVDKRFSLTDNRTAHIFTITITDLRTDDQGKYWCAVKTGIGSFDKYT
Nilt79-D1     -TGHEGRKVKIKCPYD-AKFEKMEKYLCRGKCSIG-----YKDIPVESGS-EPKDKRFSLTDNRTAHIFTITITDLRTADQGKYWCAVKTRLGKFDDYK
Nilt11-D1     VSGYRGETLDIRCSYLELSGTEYPKFFCKGECYFG-----FENVIVGSGS-PAKDQRFSVNDDKTNRVFTVTITDLRTEDAGQYWCALEDG--------
Nilt10-D1     LSGHRGQRFDIKCRYNSG-YESNVKYFCKGKCHYG-----NKEIVAHSGT-PAKDERFSLTDDTTNNVFTITITDLRTEDAGTYWCAVERVL-LPDVYS
Nilt1-D1      VSGHRGEKLDIRCSYESG-YESNSKYFCKGKCNYG-----TRNIIVESGS-PAKDERFFMNDNKESRVFTITITDLRTDDAGQYWCGVRRSLT--DVYS
Nilt16-D1     VSGHRGEKLNIRCPYESG-YESNSKYFCKGKGILG-----WKDIIIKSGS-PIKDKRFSLSDDKKSRVFTVTITDLRTEDAEQYWCAVERSLPLTDLYS
Nilt15p-D1    VSGHRGEKLDIRCPYDPG-YESNSKYFCKGWCITG-----LKNTIVESGS-PAKDERFSLNDNKTSRVFTITITDLRTEDAGQYWCAVERSLL--DDYS
Nilt18-D1     ASGHRGEKLDIRCSYDPG-YESNSKYFCKGKCTPG-----FKDIIVESGS-SPKDKRFSLTDDTTNKVFTVTITDLRTEDAGQYWCAVEKSVF--DVYS
Nilt20p-D1    ASGHRGEKLDIRCSYESG-YESNSKYFCKGKCTPG-----FKDIIVESRS-SPKDKRFSLTDDTTNKVFTVTITDLRTEDEGQYWCAVEKSVF--DVYS
Nilt17-D1     VSGHRGEKLDIRCPYKSG-YESYSKYFCKGSCFTG-----FKDTVVESGS-PAIDERFSLTDDKKNRVFTITITDLRSADAGQYWCAVQRSLLIEDLY-
Nilt22p-D1    ASGHRGEKLDIRCPYESG-YESNSKYFCKGECTPG-----FKDIIVESGS-PAKDERFSLTDDKKNRVFTITITDLRTEDEGQYWCAVRRS--------
Nilt21-D1     ASGHRGEKLDIRCSYESG-YESNSKYFCKGKCNFG-----SKNIIVESGS-PAEDDRFSLTDDTINRVFTITITDLRTEDAGQYWCAVERSLLIKDLYS
Nilt14-D1     VSGHRGEKLDIRCPYESG-YESNSKYFCKGECNFG-----NKNIIVKSGS-PAKDKRFSLTDDKKSRVFTITITDLRTEDEGQYWCAVRR---------
Nilt19-D1     ASGHRGEKLDIRCSYESG-YESNSKYFCKGECNFG-----NKNIIVESGS-SAKDKRFSLTDDKKNRVFTITITDLRTEDAGQYWCAVRRGLLI-----
Nilt9-D1      VSAPVGGSVTFECSHFLA--FLSTKYFCRDSCAEE-----NILIQSETGENPTRRGRYSLNE-KGS-DFTVTIADLRLSDSGTYVCAVERLLK--DTYT
Nilt3-D1      TQGYKGRNITITCSHKWA--SDNIKYFCRDPCKDT-----TEVLVK---SDQSPKGRYTLKD-TGD-VFTVTITNLQESDSGVYWCGVERVGL--DTYN
Nilt5-D1      TQGYKGRNITITCSHKWA--SDNIKYFCRDPCKDT-----TEVLVK---SDQSPKGRYTLKD-TGD-VFTVTITNLQESDSGVYWCGVERVGL--DTYN
Nilt4-D1      IQEYTGKYVIIICSHKWA--SDNFKYFCRDPCTYK-----DGALVS---SKQAEVGRYRLKD-LGDGSFTVTIADLQESDSGIYWCGVERSIK--DTYE
```

**Figure S4. Alignment of zebrafish NILT Ig domains with $Cx_3C$ motif.**
NILT Ig domains encoded by reference genome sequences (**Supplementary Table S2**) and possess a $Cx_3C$ motif (shaded orange). Sequences were aligned with Clustal Omega (Sievers and Higgins, 2014) and formatted with Boxshade (https://embnet.vital-it.ch/software/BOX_form.html). The cysteine pair indicative of an Ig domain is shaded red. Identical residues are shaded black and structurally similar residues are shaded gray. Sequence names refer to the NILT gene name and the position of the Ig domain within the predicted protein (e.g. D1 is the most membrane-distal Ig domain in the reported sequence). Pseudogenes are included for comparison. Note that although the Nilt16-D1 and Nilt57-D5 Ig domains lack one of the cysteines in the $Cx_3C$ motif, they are included in this group based on sequence similarity.

```
Nilt54p-D4    KTTRLGGDVNISCQTPEE----HIAYFCKEDDNDCYNRIRQNFSSA------EVPQ-----INPAEFVSISNVSVRDAGVYWCGAETRDTHLT---------
Nilt75-D4     KTTRLGGEVNISCQTPEE----HIAYFCKEDDNDCYNRIRQNFSSA------EEPQ-----MNPAEFVSISNVSVRDAGVYWCGAETRDTHLT---------
Nilt37p-D2    --------------------------FCKEEYNKCSQIRRA----S------KEPQ-----MTPGER--VYNVSVRDAGVYWCGAETRDTHLT---------
Nilt40-D2     KSVHLGGDVNISCQMAKK----MKVYFCKEDNQRC-QIMRA----S------KEPQ-----MTPGER--VYNVSVRDAGVYWCGAETRDTHLT---------
Nilt55-D1     VTGVAGGSVIFNFQYKDKIS---VERFCRDADDECIQAQ------SHLWLPEEKLTMTDNKSVNYLSVLIRNLTVNDSGTYEWKDNKD--------------
Nilt76p-D1    VTGVAGGSVIFNLLYKDKMTEESVKSFCRVSDDKCTKAR------NNLYLPEEKFTMTENKSINFLSVLIRNLTINDRGMYRWKANNKGS------------
Nilt66-D1     -SSFAGAALMIKCEH-PQ-HQTKTRYICKESSAGC----------SGEWRKNGDVSVDEDTRAGVLMVFFRELKAADAGTYRCGVKDSEYTERFTQLQLK--
Nilt49p-D1    VTGSSGAALMIKCEH-PQ-HKTNTRYICKESSAGC----------SEEWMENGDVSVDDDTRAGVLMVFFRELKAADAGTYRCGVKDSEYTERFTQLQLK--
Nilt37p-D1    VIGYSGAALMIKCEH-PQ-HKTKTRYICKESSAEC----------SEEWRKNGDVSVDVDSRAGVLMVFFRELKAADAGTYRCGVKDSEHTERFTQLQLK--
Nilt75-D1     VIGYSGAALMIKCKH-PQ-HKTKTRYICKESSAGC----------SEEWRKNGDVSVDDDSRAGVLMVFFRELKAADAGTYRCGVKDSEYTERFTQLQ----
Nilt54p-D1    VIGSSGAALMIKCEH-PQ-HKNKTRYICKESSAGF----------SEEWMKNGDVSVDEDSRAGVLMVFFRELKAADAGTYRCGVKDSEYTERFTQLQLK--
Nilt40-D1     VIGYSGAALMIKCEH-PQ-HKTKTRYICKESSAGC----------SGEWMKNGDVSVDEDTRAGVLMVFFRELKAADAGTYRCGVKDSEYTERFTQL-----
Nilt75-D3     VIGSSGATLMIKCEH-PQ-HKTNTKYICKESSAGC----------SEEWRKNGDVSVDDDSRAGVLMVFFRELKAADAGTYRCGVKDSEYTERFTQLQLK--
Nilt54p-D3    VIGYSGAAVMIKCEH-PQ-HKNKTKYICKESSAGC----------SGEWMKNGDVSVDEDSRAGVLMVFFRELKAADAGTYRCGVKDSEYTERFTQLQLK--
Nilt70-D3     VIGYSGAALMIKCEH-PQ-QKTNTKYICKESSAGC----------SEEWRKNGDVSVDEDTRAGVLMVFFRELKAADAGTYRCGVKDSEYTERFTQLQ----
Nilt34p-D1    VIGYSGAALMIKCEH-PQ-HKTNNKYICKESSAGC----------SGEWRKNGDVSVDEDSRAGVLMVFFRELKAADAGTYRCGVKDSEYTERFTQLQLKVR
Nilt43p-D1    VIGYSGAALMIKCEH-PQ-HKTNTKYICKESSAGC----------SEEWMKNGDVSVDEDSRAGVLMVFFRELKAADAGTYRCGVKDSEYTERFTQLQLK--
Nilt26-D2     VRRHSGESITIDYTYEEK-HKSREKSVCKIGEYLCETLISISR--PAEKKDSGRFSIHDDRSAGLLRVFIRELNVQDSGEYRIKVRHSEDYSFFSEFELV--
Nilt63-D2     VRRHSGGNITIDYTYEKQ-HKSREKSVCKIGEYQCETLISISR--PAE-KDSGRFSIHDDRSAGLLHVFIRELNVQDSGEYRIIVRHSEDYSFFSEFDL---
Nilt62p-D1    VSGYSRGNVTIDYIYGKQ-HKNHQKYVCQSANHSCFALISING--SAEWKHRGQFSVHDDRSADLLRVFIRELKIQDSGDYRIIVRASED------------
Nilt60p-D1    -RGYSGGNININYSYEIQ-HRNHQKSFCK-GIYKCITLISING--SAESEDSGRFSIHDDRSAGLLRVFIRELNIQDSGGYRITVRVSEDYSFFS-------
Nilt27-D2     VRGYSGRNIIIKYRYDKQ-HKNCQKYVCKTGENHCLSLISIYG--GAEWEHSGRFSIRDNRSAGLLRVFIRELNVQDSGRYRIIVRHSDYYS----------
Nilt28-D2     VRGYSGGHVIINYSYAIQ-HKNSQKHVCKTGENLCLSLISISG--GAEWEHSGRFSIHDNRSAGLLRVFIRELNVQDSGEYRIIVRASED------------
Nilt67-D2     VRGYSGGHVIINYSYAIQ-HKNSQKHVCKTGENHCLSLISISG--AAEWKHSGRFSIHDDRSAGLLRVFIRELNVQDSGEYRIIVRASED------------
Nilt29-D2     VRGYSGGHVIINSSYEIQ-HKNSQKHVCKTGENHCLSLININA--AAEWKDSGRFSIHDDRSADLLRVFIRELNVQDSGEYRIIVRESEDYS----------
Nilt68-D2     VRGYSGRNIIINYSYEIQ-HKNSQKYVCKTGENHCLSLININA--AAEWKDSGRFSIHDDRSAGLLRVFIRELNVQDSGGYRIIVRESEDYSL---------
Nilt27-D1     VTGYLGGGVTITCKYDEQ-YKTNAKYFCKGKGLSCFELIKTDTKSENKWVQSGRFSLYDNTTAAVFTVTIRNLTRQIYGTYHCGVEVYG-------------
Nilt26-D1     -TGY--SGVTITCKYDEE-YETNEKYFCKGKWSECEDQIRTEM--KHHWIRSGRFSLFDDTTAAVFTVTIRNLRKQDSGIYHCGIKRFAFDH-GTKVN----
Nilt63-D1     VTGYSGGGVTIRCKYDEE-YETNAKYFCRGEWPECTDQIRTEM--KHQWIRSGRFSLFDDTRAAVFTVTIRNLRKQDSGMYYCGIKIFGPDH-DTKVNLK--
Nilt25-D1     VTGYSGGGVTITCKYDEG-YETYVKYFCKGQWSECTDRIRTGT--KDTWVQSGRFSLYDDTTAAVFTVTIRDLTEEDSDMYYCGTDIYGKVDSATEVNLK--
Nilt61-D1     VTGYSKGGVTITCKYDKG-YETYVKYFCKGEWSGCTDQIKTAT--QDKWIKSGRFSLYDNTKEAHLIVTIKDLREQDSDIYYCGTEVYGYDL-STGVNLK--
```

#### Figure S5. Alignment of zebrafish NILT Ig domains with $Cx_6C$ motif.

NILT Ig domains encoded by reference genome sequences (**Supplementary Table S2**) and possess a $Cx_6C$ motif (shaded orange). Sequences were aligned with Clustal Omega (Sievers and Higgins, 2014) and formatted with Boxshade (https://embnet.vital-it.ch/software/BOX_form.html). The cysteine pair indicative of an Ig domain is shaded red. Identical residues are shaded black and structurally similar residues are shaded gray. Sequence names refer to the NILT gene name and the position of the Ig domain within the predicted protein (e.g. D1 is the most membrane-distal Ig domain in the reported sequence). Pseudogenes are included for comparison. Note that although the Nilt54p-D1 Ig domain is lacking one of the cysteines in the $Cx_3C$ motif, it is included in this group based on sequence similarity.

```
Nilt57-D4     EAAFIGESVKLTCIHREKN--TVNHICKENHDKICQNIVSSNK------GRVQ----ISDSTDGVFTVSISNVSVRDAGVYWCGAKTRDTHL-TFISL
Nilt75-D2     -SVQLDGEVKISCQILEEQKVHFSYFCKEDDHKKCQKTRASEE------PQRTPAG-SVGSEERVFTVSISNVSVRDAGVYWCGAETRDTHL-T----
Nilt66-D2     ESVQLGGEVKISCQIPDEQK---VHFCKEDDHKSCQIMRASED------PQLTPAG-SVGNKERVFTVSISNVSVRDAGVYWCGAETRDTHL-T----
Nilt50p-D1    ESVHLGGEVKISCQIPEQQNK--NYFCKEDDHKSCQIRRVSKD------PQLTPAG-SVGSEERVFTVSISNPSVRDAGVYWCGAETRDTHL-T----
Nilt70-D4     ESVHLGGEVKISCQIPEEQNK--NYFCKEDDHKSCQIRRASED------PQLTPAG-SVGSEERVFTVSISNVSVRDAGVYWCGAETRDTHL-T----
Nilt44p-D1    ESVHLGGEVKISCQIIEEQK---VYFYKEEYNQSCQIRRASED------PQLTPAE-SVGSEERVFTVIISNASVRDAGVYWCGAETRDTHL-T----
Nilt54p-D2    ESVHLGGEVKISCQIPEEQK---VLFCKEEYNQSCQKTRASED------PQLTRAG-SVGGEERVFTVSISNVSVRDAGVYWCGAETRDTHL-T----
Nilt24-D2     LSAAAGGSVNISCRYPQSHTADVKFICRRSGADLCAEETRSEE-ESGGRKAEGKIQLYDDREEQLLTATISHESPEHSAEYWCGVQSAGHK-------
Nilt25-D2     LSAAAGGSVNISCRYPQSHTADVKFICRRSGADLCAEETRSEE-ESGGRKAEGKIQLYDDREEQLLTATISHESPEHSAEYWCGVQSAGHKS------
Nilt58-D1     VTGYSGGVTITCKYDRQYKKNTKYFCKGQKFLSCSKLIKTEAKSDNKWVQKDRYSLFHNTTAADFTVTIRNLTVWDSGTYYCGVEVYGLDPSTKVNL
Nilt23-D1     VTGYSGGVTITCKYAAQYKTKAKYFCKGQWLSTCSDQIKTEI--KNKWVQSGRFSLYDNTNAAVFNVTIRNLTEQDSGTYYCGVDLQISDIYTKVNL
Nilt68-D1     VIGYSGGVMITCKYDEQYKTSAKYFCKEQWISPCSELIR--INSEEKWVQSGRFSLFDNRTAAGLTVTIRNLTEQDSGTYHCGVETYGPDPGTKVNL
Nilt28-D1     VIGYSGGVMITCKYDKGFETYPKYFCKEQWISPCSELIRTEINSEEKWVQSGRFSLIDNKTASGLTVTIRDLTEQDSGTYYCGVEIYGLDPGTKVNL
Nilt29-D1     VIGYSGGVMITCKYDRGFETYPKYFCKEQWISPCSELIRTEINSEEKWVQSRRFSLIDNTTAAFLTVTIRDLTEQDSGTYYCGVETYGPDLGTKVNL
Nilt67-D1     VIGYSGGVMITCKYDRGFETYPKYFCKEQWISPCSELIRTEINSEEKWVQSGRFSLIDNKTAAGLTVTIRDLTEQDSGTYYCGVETYGPDLGTKVNL
Nilt46p-D1    LTKYEGDDVSIQCR---HHDGDQKSFCRAHEASTCVKDG-----DSLETIRDARFSISDEASAGVFTVNITDIRADDSGVYWCGAHV-----------
Nilt37p-D4    LTKYEGDDVSIQCR---HHDGDQKSFCRAHEASACVKDG-----DSLETIRDARFSISDEASAGVFNVNITDIRADDSGVYWCGAHA-----------
Nilt75-D6     LTKYEGDDVSIQCR---HHDGDQKSFCRAHEASACVKDG-----DSLETIRDARFSISDEASAGVFTVNITDIRADDSGVYWCGAHVITKVNLT----
Nilt69-D4     LTKYEGDDVSIQCR---HHDGDQKSFCRAHEASACVKDG-----DSLETIRDARFSISDEASAGVFTVNITDIRADDTGVYWCGAHAV----------
Nilt66-D4     LTKYEGDDVSIQCR---HHDGDQKSFCRAHEASACVKDG-----DSLETIRDARFSISDEASAGVFTVNITDIRADDSGVYWCGAHLITKVNLT----
Nilt40-D4     LTKYEGDDVSIQCR---HHDGDQKSFCRAHEASACVKDG-----DSLETIRDARFSISDEASAGVFTVNITDIRADDSGVYWCGAHLI----------
Nilt30-D4     LTKYEGDDVSIQCR---HHDGDQKSFCRAHEASACVKDG-----DSLETIRDARFSISDEASAGVFTVNITDIRADDSGVYWCGAHAV----------
Nilt54p-D6    LTKYEGDDVSIQCR---HHDGDQKSFCRAHEASACVKDG-----DSLETIRDARFSISDEASAGVFTVN-----------------------------
Nilt57-D6     --ASVRGSASISCKCIRKHTQHQRFFCKGNQPNICVHAGVL---VSFNKPANSRFSLTDECSAGVFTVNISQLTEEDSGKYWCGEESSGSFLFTEVHL
Nilt78-D2     VSGEPGKHLSITCSYTRDLKELIRFLCKGSNPSHCKNII-K---VSSETKTNGRFSLTDDS-ERNFTVNISDLTEEDSGIYWCGAEDNQTQEY-----
Nilt79-D2     VSGEPGKHLNITCSYTRDLKELIRFLCKGSDPSDCKKMI-K---VSSETNTTGRFSLTDDS-ERNFTVNIRDLTEEDSGIYWCGAEEKQTR-------
Nilt2-D2      VSGEPGKHLSITCSYTRDLKEHIRFLCKGSDPSDCKNMI-K---VSSETKTNGRFSLTDDS-ERVFTVNIRDLAEGDSGIYWCGAEEKQTQEY-----
Nilt56-D2     VSGEPGKHLSITCSYTRDLKELIRFLCKGSDPSDCKNII-K---VSSKTKTNGRFSLTDDS-ETVFTVNIRDLTEEDSGIYWCGAAEKQAQEY-----
```

#### Figure S6. Alignment of zebrafish NILT Ig domains with $Cx_7C$ motif.

NILT Ig domains encoded by reference genome sequences (**Supplementary Table S2**) and possess a $Cx_7C$ motif (shaded orange). Sequences were aligned with Clustal Omega (Sievers and Higgins, 2014) and formatted with Boxshade (https://embnet.vital-it.ch/software/BOX_form.html). The cysteine pair indicative of an Ig domain is shaded red. Identical residues are shaded black and structurally similar residues are shaded gray. Sequence names refer to the NILT gene name and the position of the Ig domain within the predicted protein (e.g. D1 is the most membrane-distal Ig domain in the reported sequence). Pseudogenes are included for comparison. Note that although the Nilt44p-D1 Ig domain is lacking one of the cysteines in the $Cx_3C$ motif, it is included in this group based on sequence similarity.

```
Nilt57-D1  VHGCTGGSLLFKCMYSNQKSSDVENDGAYFSR-D-----NPYKM--VINTKIQSCWSHNGRFAVYDDE--IRKFILVLIKNLSREDDGTYTCGHDHKWNHNVELV--
Nilt35p-D1 VTGYSGGSLMLDSGRS-----SSSNSLYLLKHTHGGWG----LI--ILENSRQDLWVSDGKFTLYLNK--NRN--MLFIRDLTPQDSGRYTI--------------
Nilt38p-D1 VTGYSGGSLMLDSGRS-----SSSNSLYLLKHTHGGWG----LI--ILENSRQDLWVSDGKFTLYLNK--NRN--MLFIRDLTPQDSGRYTVGLS-----------
Nilt73p-D1 VTGYSGGNLLLDSRKP-----WLSDFAKYMAK-LQEWRPAKWRT--IIHNKKHLKWINEGRFTLFSGN--NGY-LMIFIRDLNLSDEGEYRIRVQDQY---------
Nilt64p-D1 VTGYSGGSLILDSGKL-----WFSESAKYMVK-HNPW-----RT--IIDYKKHSKWTAEGRFTLFCNT--HGN-LVIFIRDLKSQDAGKYRVRVLD-----------
Nilt31p-D1 VTGYSGRSLMLNSGRL-----WHKNYAKYMHK-LPEW-----KT--LMYYKIHEKWINEGRFTLFENS--DGN-LMIFIRDLHTDDAGVYRIGVYDD----------
Nilt70-D1  -----------DSGKR-----WFSDSAKYLAK-LPEW-----RT--IINDRKHERWINEGRFTLFKNT--DAN-LMIFIRDLHTDDAGRYAVDVEQGNRIYIILNVG
Nilt47p-D1 -TGYSGGSLMLDSGKL-----WISDSAKYLAK-LPDW-----RI--IINDTKHERWINEGRFTLFKNT--DEN-LMIFIRDLNTDDAGRYAVDVDQE----------
Nilt51-D1  -TGYSGGSLMLDSGKL-----WVSDSAKYLAK-LPDW-----RT--IINDTKHERWINEGRFTLFKNN--DGN-LMIFIRDLNTDDAGRYAVDVEQG----------
Nilt71p-D1 -TGYSGGSLMLDSGKL-----WISDSAKYLAK-LPDW-----RT--IINDRKHERWINEGRFTLFKNT--DGN-LMIFIRDLNTDDAGRYAVDVEQG----------
Nilt30-D1  -TGYSGGSLMLDSGKL-----WISDSAKYLAK-LPNW-----RT--IINETKHERWINEGRFTLFRNN--NGN-LMIFIRDLHTDDAGEYWLG--------------
Nilt69-D1  -TGYSGGSLMLDSGKL-----WFSDSAKYLAK-LPNW-----MT--IINDRKHERWINEGRFTLFRNN--DGN-LMIFIRDLHTDDAGEYWLG--------------
Nilt41p-D1 -TGYSGRSLLVSSWKP-----WHENDTKFMGK-LQD------RP--IIKNRKHHRWINEGRFTLFCNN--LAS-LVLYIKELNKQDVGKYWIRVQGKW---------
Nilt52p-D1 VTGYLGGNLLVSSLEP-----WHESDAKFMEK-LRG------SI--IINNRKQRQWINEGRFTLFYTD--YRN-LVIYIKELVTEDAGTYRIKVKGKW---------
Nilt57-D2  --SYAGQETTIICKYQD----KFKKSTKVFYTLNGDPVHMLNSS--------S--QSSVKRFSLSDSH---KDHFNVTISSVSAEDDGDYLCGVERSDDQRSSSI--
Nilt74p-D1 VMVKSGETASFNCKYSQ----SRFNS-EVIFKAQQSSIEELKYT--------R--WNSGAKFTISNDR--EKNLFSVKITAVNSFDGAVYLCGIHQYHQLYSY----
Nilt36p-D1 QEVDLRRTGNFICEFSH----NHINDEKLVFKEGKNSIDMILNS--------T--WRMSTRFGLINMI--DKQHFKVRINAVTPDDGGVYLCGVWVNKHSYS-----
Nilt39p-D1 QEVNLGRTGIFFCEYSQ----NHINHDKVVFKEGKNSIDEIINS--------T--WKQ-VRFSWFNIR--DKRHFNVIITAVKADDGGVYLCGVWINEHSYS-----
Nilt42p-D1 VMVNNGETASFNCESSQ----SQINNHKIILKKAKDYIELIYNS-----------WYNEGRVSVSDDRPDQKKLFSVRITAVTPEDGGVYFCGVQFYRNSH------
Nilt65p-D1 VKVNSGETAVFSCDYSR----NHIYDAKVLFKERQNLIEEVIHS--------TGT-RNKERFSISDDR--RNNFFNVRISTVKPDDGGVYLCGVSISRYSS---IIG
Nilt33p-D1 LVVKSREAAAFTCEYSL----NQTSGGKTIFKTDNDIFDEVIST--------TYTWDRKERVSISDDR--QRKLLSVSITAVTADDGGVYLCGVWVKDNSYSYYITN
Nilt48-D1  VTVNSGETAVFSCEYSH----NQIHHAKVIFKEERDSVKSVIHT--------TARIREEGRVHMSDDG--QRNVLSVSITAVTADDGGVYLCGVWVRENSYNYSTLH
Nilt70-D2  VTVNSGETAVFSCEYSH----NQIHHAKVIFKEERDSVKSVIHT--------TARIREEGRVHMSDDR--QRNVLSVSITAVTADDGGVYLCGVWVRENSYNYSTLH
Nilt53p-D1 VMVNNGETNSFSCKYSH----NQIHNFKAIFKEGKNSIEVIHNT-----------WKKTDRFSVSDDG--QRNSLHVSITAVTADDGGVYLCGVWVNKHSYSYYITN
Nilt30-D2  VIMTVGETGSFSCDYSH----NHINDPKIIFKEEQHSIKMIYST-----------LNKKDRFNISDDG--LRNSFSVSITAVTADDGGVYLCGVWVNKHSYS-----
Nilt69-D2  VIMTVGETGSFSCDYFH----YHINDPKIIFKEEQHSIKMIYST-----------LNKKDRLNISDDG--LRNSFSVSITAVTADDGGVYLCGVWVNKHSYSYYITN
Nilt32p-D1 VMVHSGESGSFSCEYSH----NHINDEKVVFKEGRDSIEMIYST-----------LNKKERFSISDDR--QRKLFNVSITAVTTDDGGVYLCGVWVNKHSYS-----
Nilt51-D2  VTVNSGETAVFSCEYSH----NQINHAKVILKPERDSIEMIYST-----------LNKQDRFSISDDR--QRNSFSVSITAVTAGDGGVYLCGVWVNKHSYSYYITN
Nilt72p-D1 VTVNSGETAVFSCEYSH----NQIHHAKVILKPGRDSIEMIYST-----------LNKQDRFSISDDK--QRNSLSVSITAVTADDGGVYLCGVWVNKHSYS-----
Nilt55-D2  QTAYVQGTAVIHCRYPK----TNHDYSKQLFKEQNGSLEA-IFTSR---------EASDPRFSINLNH-GEEHFFTVTLTNVSRHDEGLYFCAAHGKMSIRYSLLFT
Nilt77p-D1 QTAYELGIFVIHCKYPS----TLHDFPKQLFKEQNGSLQA-IFTYGSVFSSKQTKISIDPRFSMNVNH-SEEHLFTVTLINVSRHDEGLYFCAAHGVMTISYSLLFT
Nilt6p-D1  ITAAEGGEAVFNCPYYE----GYENSKKHVYKEPYKTNSP------ILQADGGKS-VSNGRFTLKDDP--QARLFTVTIRDLQMNDAGLYGCAPGL-----------
Nilt12-D1  VTAAEGGEAVINCPYGG----GYETSYKYFYIGPYKDSTL------WLQSYGGESPVFNGRFTLKDDH--KARLFIVTIRDLQMNDAGEYNCAAGWRD-------SR
Nilt13-D1  VTAAEGGEAVINCPYGE----GYETSYKYFYKGPYRESTL------MLKSDGGQSPVFSGRFTLKDDH--KARLFTVTIRDLQMSDAGEYTCTAGWGD-------SR
Nilt57-D3  VQAYIGRIVFIKCKFPQ----KIKENKKFFMRAESQ----------KIMLDEQNQWIHHDNVHMYDNT--TEGHLKVFISDLSAANEGTYRCGVNID-Q---DDLYT
Nilt7-D1   -TEYEGKSITFRTSYTE----GFETNSKYFSRSDT------FFSEKLVETSESNRWTTHGRFAIFDNT--SAHVIIATILNLTMEDSGSYTVGVDVTLL---PDYDA
Nilt8p-D1  -TEYEGKSITFSTSYLE----SLETNSKYFSRSG-------YFCEKLVETSESNRWTTHGRFAIFDNT--STHLITTTILNLTVEDSGLYIVAVDVTLW---PDPET
Nilt24-D1  VTGYSGGGVTITCKYDE----QYKTNSKYFCKGKYIPT-----GPDLIKTEIKNKWVKKDKYFLYDNT--TAAVFTVTIINLTEKDSGTYYCGVDIHYH---TDIYT
Nilt59p-D1 VMGYLGGGVMITCNYDR----VYETNAKYFCKGKKPAIFPIQWCYDLIKTEINSKWIQKDRFSLYDDT--AAAVFTVTIRNLTKQDSEIYQCAVDTS-G---IDQYT
```

#### Figure S7. Alignment of atypical zebrafish NILT Ig domains.

NILT Ig domains encoded by reference genome sequences (**Supplementary Table S2**) and classified as atypical. Sequences were aligned with Clustal Omega (Sievers and Higgins, 2014) and formatted with Boxshade (https://embnet.vital-it.ch/software/BOX_form.html). The cysteine pair indicative of an Ig domain is shaded red. Identical residues are shaded black and structurally similar residues are shaded gray. Sequence names refer to the NILT gene name and the position of the Ig domain within the predicted protein (e.g. D1 is the most membrane-distal Ig domain in the reported sequence). Pseudogenes are included for comparison.

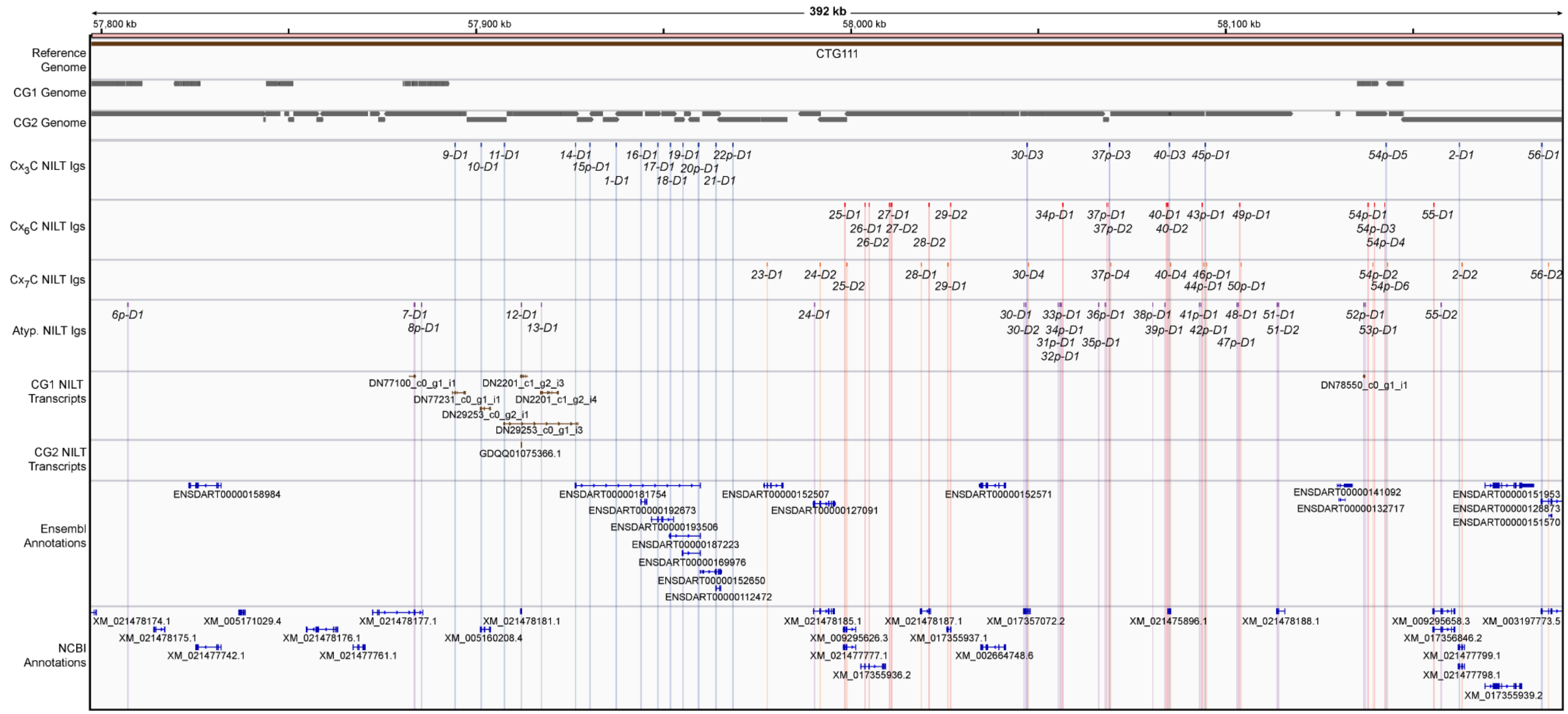

**Figure S8. Integrative Genomics Viewer - Region 1**.
Annotations were mapped to GRCz11-Chr1 modified to include Zv9-scaffold100 between adjacent scaffolds CTG104 and CTG111 as described in the main text (Figure 4). Genomic coordinates 57,797,367 - 58,189,758 bp are shown with transcript names for Ensembl and NCBI gene annotations. Pseudogenes, ncRNA, and miscellaneous RNA are not displayed. The image from IGV v2.8.10 (Robinson et al. 2011) was exported and manually edited in Adobe Illustrator.

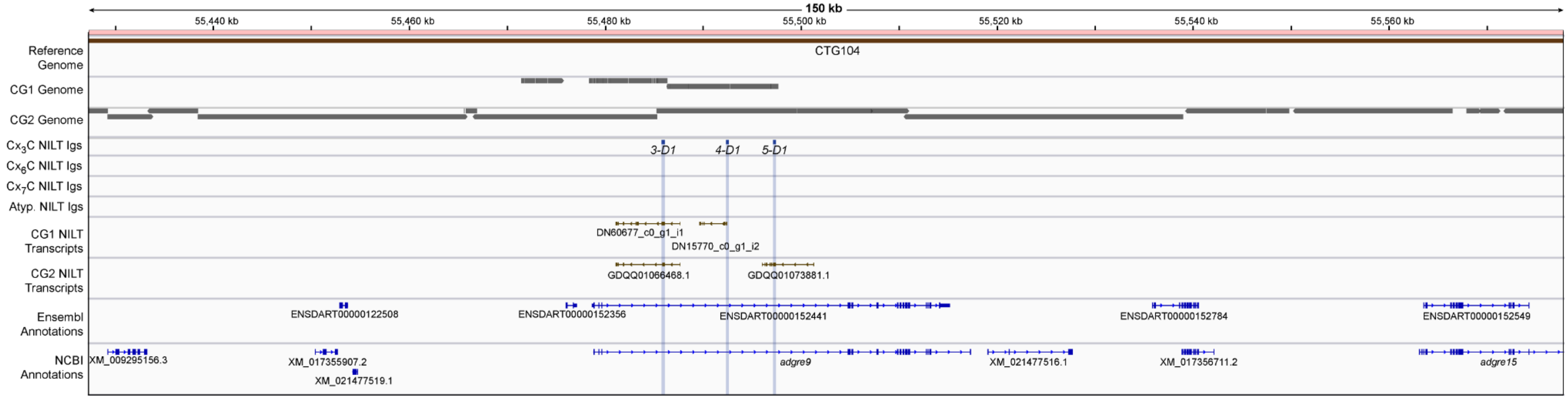

**Figure S9. Integrative Genomics Viewer - Region 2.**
Annotations were mapped to GRCz11-Chr1 modified to include Zv9-scaffold100 between adjacent scaffolds CTG104 and CTG111 as described in the main text (Figure 4). Genomic coordinates 55,427,288 - 55,577,739 bp are shown with transcript names for Ensembl and NCBI gene annotations. Pseudogenes, ncRNA, and miscellaneous RNA are not displayed. The image from IGV v2.8.10 (Robinson et al. 2011) was exported and manually edited in Adobe Illustrator.

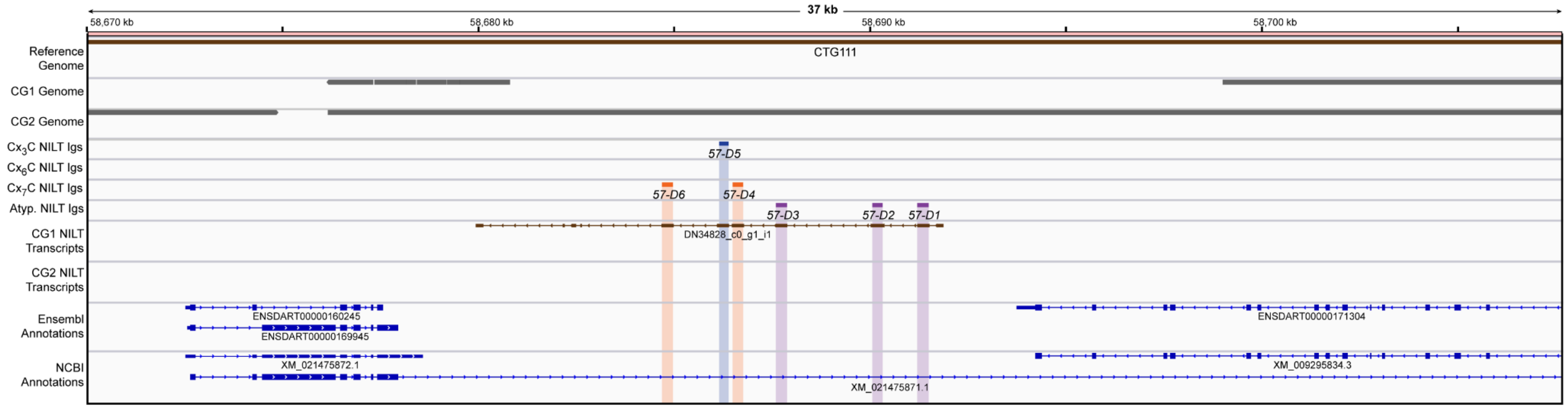

**Figure S10. Integrative Genomics Viewer - Region 3.**
Annotations were mapped to GRCz11-Chr1 modified to include Zv9-scaffold100 between adjacent scaffolds CTG104 and CTG111 as described in the main text (Figure 4). Genomic coordinates 58,670,034 - 58,707,646 bp are shown with transcript names for Ensembl and NCBI gene annotations. Pseudogenes, ncRNA, and miscellaneous RNA are not displayed. The image from IGV v2.8.10 (Robinson et al. 2011) was exported and manually edited in Adobe Illustrator.

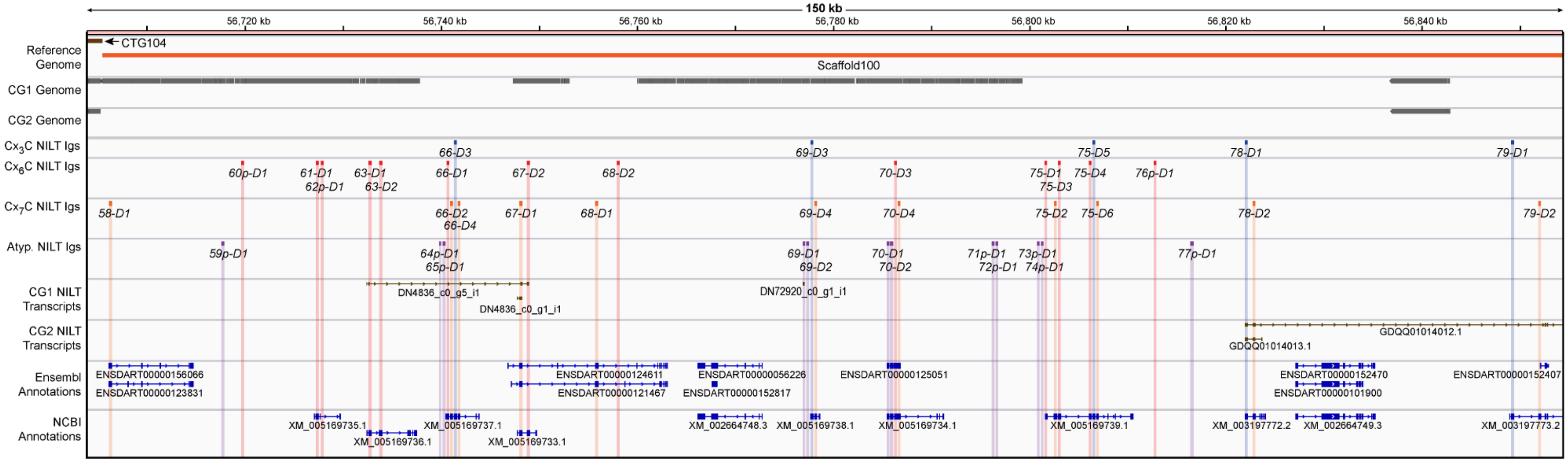

**Figure S11. Integrative Genomics Viewer - Region 4.**
Annotations were mapped to GRCz11-Chr1 modified to include Zv9-scaffold100 between adjacent scaffolds CTG104 and CTG111 as described in the main text (Figure 4). Genomic coordinates 56,703,879 - 56,854,330 bp are shown with transcript names for Ensembl and NCBI gene annotations. Pseudogenes, ncRNA, and miscellaneous RNA are not displayed. The image from IGV v2.8.10 (Robinson et al. 2011) was exported and manually edited in Adobe Illustrator.

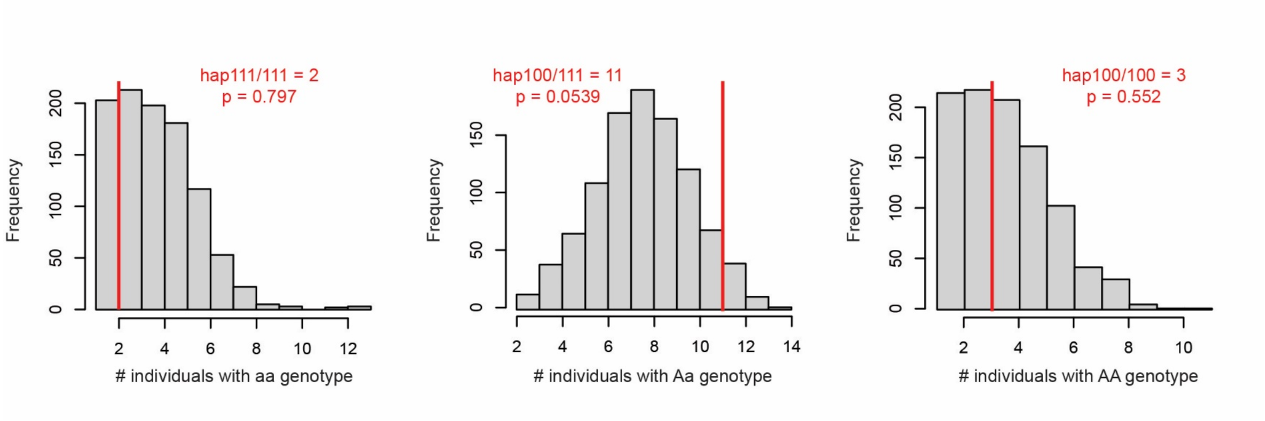

**Figure S12. Statistical analyses of haplotype distribution.**
As the sample size (n=16) of individuals genotyped for NILT haplotypes was low relative to the total population, we assessed if observed genotype frequencies are within the expectations for Mendelian ratios under stochastic sampling using Monte Carlo simulation. We simulated a population of one hundred thousand individuals with a frequency of genotypes at the classic ratio of 25% homozygous aa, 50% heterozygous Aa, and 25% homozygous AA. Sixteen individuals were then randomly sampled from this population one thousand times, and the frequency distribution of each genotype was compared to our empirically observed frequencies to assess significance. Simulations were conducted in R using code modified from Dornburg et al. (2016). The gray histogram represents the distribution from the simulation and the red line indicates the observed frequencies.

>Nilt1(BN001234)
MLVILLLVSSFSTALYDHIWVSAVVGAPRSVSGHRGEKLDIRCSYESGYESNSKYFCKGKCNYGTRNIIVESGSPAKDERFFMNDNKESRVFTITITDLRTDD
AGQYWCGVRRSLTDVYSKVLLNVKQNQENTEGSSVSSFSNTPSYFSSTEVNLQSSSSTDQHNSTASHTDLGSVVGGLGSVLLVLLLCSGTFLILKKRKRKCET
ALPLQNVQQNTENGCVYENDLYGDVIIATKPSSNQKAASDLHTHSVITVYATVTN-

>Nilt2(BN001235)
MIHISEDKLLIFTLLILMAVVDSEMKTFTGHEGGKVEIQCSYAESEYEENQKYLCRGECSINPFPFNIKDIPVESGSEPKDKRFSLTDDTTAHIFTITITDLR
TADQGKYSCAVKTRLVKLDDYKEIYLEIKQVSRVSGEPGKHLSITCSYTRDLKEHIRFLCKGSDPSDCKNMIKVSSETKTNGRFSLTDDSERVFTVNIRDLAE
GDSGIYWCGAEEKQTQEYTWISAIDLHITEATSERRLPKNTSTASFHTSKPADPASSRPVTPSSPASSSSTSMLSSWKRLLCTVFIILITSVMLTVFGLSVFI
FLRWKPQTKGGGNAAHDLTEKLLNTEEAPGDIYTVCHYEELIHTDDHPAGLGSGRPVFGEQDPLTCSTIIFNTGLHSDTTSD-

>Nilt2(XM_021477798.1)
MIHISEDKLLIFTLLILMAVVDSEMKTFTGHEGGKVEIQCSYAESEYEENQKYLCRGECSINPFPFNIKDIPVESGSEPKDKRFSLTDDTTAHIFTITITDLR
TADQGKYSCAVKTRLVKLDDYKEIYLEIKQVSRVSGEPGKHLSITCSYTRDLKEHIRFLCKGSDPSDCKNMIKVSSETKTNGRFSLTDDSERVFTVNIRDLAE
GDSGIYWCGAEEKQTQEYTWISAIDLHITEATSERRLPKNTSTASFHTSKPADPASSRPVTPSSPASSSSTSMLSSWKRLCMINLVVFLRIMTF

>Nilt3(DN60677_c0_g1_i1)
MRFVCVLWLWTFISEFKTSTTDEVSTQGYKGRNITITCSHKWASDNIKYFCRDPCKDTTEVLVKSDQSPKGRYTLKDTGDVFTVTITNLQESDSGVYWCGVER
VGLDTYNKVTLTVSEDNTWVPEEVIHVRTTEPQISLQTRAEIHSTTPRLSVSESSATKTIALFLTSTGFANATGPDDILTVPVLYIIYGGAGLVIMVIICTTG
LITVCQCRKRIKRSRSSTARSIYNGSEEASGVYENAPEDRAYACVKKSSDISKKKRQSDPIYQNIHFDTSHKDAIYGNVY-

>Nilt4(DN15770_c0_g1_i2)
iatsadiniqeytgkyviiicshkwasdnfkyfcrdpctykdgalvsskqaevgryrlkdlgdgsftvtiadlqeSDSGIYWCGVERSIKDTYEKVILEVSKD
PNESIATYTQAYTDMHTFMDSHTTYTQPELSSESPSTTASSTCRYSTLDSLTFPPSVSASVHTSPYAAVGLGLAVIILLVGLITVHLYRKRVKTSETATSADR
SDGFKEEGIQRSTEIKSTDTYEVMRSHSYSIYENLDEGQVHEDYGNL-

>Nilt5(GDQH01035306.1)
MKLICVLWIWIGLSATDGSSLHGDAGNYINISCSHSWASNNKKFFCRDPCNDKNDILVNSDKSSKGRYTLNDFGKGVFIVTITDLKESDSGVYWCGVERSIKN
TFVEVKLAVYKGKTLKQPKIPKTVTDPQSSTKSVISSSSTLDDFSSSSTAEFKYPAFSSRPAFYENTKQLDYQMFTPIILTLVVIIFVVVLCLVYMRKRKSHN
SDADAEMQSTTYTTETSLNQQAAGDYEEITETPQQISTHSTYCNVIKTHTANQNQDPPPYSLLSFQPSCTKQSPNNSAD

>Nilt9(GDQH01032966.1)
MKPLLNMRSFLVVLSSIFKVASSNSVSAPVGGSVTFECSHFLAFLSTKYFCRDSCAEENILIQSETGENPTRRGRYSLNEKGSDFTVTIADLRLSDSGTYVCA
VERLLKDTYTYLTLHVTEVPAGVTRPTTTRTFTDKTENKETLEMRPSTPPSSPETMRRSALSDPLLYVGAGLGVLLLIFTAAFFILIRLKYKQSRCFSSVISP
THEPEYANYTTGNFTAQPDSLNYSSVVFLKKTDCKVENSENNPAETHYTSVQSKTTDQDSVIYSTVSAV

>Nilt9(DN77231_c0_g1_i1)
MKPLLNMWRFLVVLSSIFKVASLNSVSAPVGGSVTFECSHFLAFLSTKYFCRDSCAEENILIQSETGENPTRRGRYSLNEKGSDFTVTIADLRLSDSGTYVCA
VERLLKDTYTYLTLHVTEVPAGVTRPTTTRTYTDKTENKETLEMRPSTPHSSPETMRRSALSDPLLYVGAGLGVLLLIFTAAFFILIRLKYKQSRCFSSVTSP
THEPEYANYTTANFTAQPDSLNYSSVVFLKKTECKVENSENNPAETHYTSVQSKTTDQDSVIYSTVSAV

>Nilt10(XM_005160208.4)
MWSFLPLLWSWISIAGVASVPDKLSGHRGQRFDIKCRYNSGYESNVKYFCKGKCHYGNKEIVAHSGTPAKDERFSLTDDTTNNVFTITITDLRTEDAGTYWCA
VERVLLPDVYSEIELLVEQEDAQTPLPQRSRSDSDSIIAIVSSGVSVLLLTGAVLFTVIIQKKKKTCGPVSSTTEALDLTLRNGEENEYEKENPTVLPNTHTA
QSVNRSAENSLPRAENTHVFYINTP**DQMYTELNTRQTDLYHSLTTD**SAQHESIYHSI

>Nilt10(DN29253_c0_g2_i1)
MWSFLPLLWSWISIAGVASVPDKLSGHRGQRFDIKCRYNSGYESNVKYFCKGKCHYGNKEIIAHSGTPAKDERFSLTDDTTNNIFTITITDLRTEDAGTYWCA
VERVLLPDVYSEIELLVEQEDAQTPLPQRSRSDSDSIIAIVSSGVSVLLLTGAVLFTVIIQKKKKTCGPVSSTTEALDLTLRNGEDNEYEKENPTVLPNPHTA
QSVNRSAENSLPRAENTHVFYINTP**DQMYTELNTRQTDLYHSLTTD**SAQHESIYHSI

>Nilt11(DN29253_c0_g1_i3)
MCVSLLLFSSICTAVVGAPVTVSGYRGETLDIRCSYLELSGTEYPKFFCKGECYFGFENVIVGSGSPAKDQRFSVNDDKTNRVFTVTITDLRTEDAGQYWCAV
RRSFTDMALTDVFSEIVLMVEQDQENTEGSSVSSFSNTPSYFSSTEVNLQSSSSTDQHNSTAHSRSEVHLCVKD

>Nilt12(XM_021478181.1)
MKFALRVCVFLLMARASVCADITVTAAEGGEAVINCPYGGGYETSYKYFYIGPYKDSTLWLQSYGGESPVFNGRFTLKDDHKARLFIVTIRDLQMNDAGEYNC
AAGWRDSRSIQLNVIRGV

>Nilt13(DN2201_c1_g2_i3)
MKFALRVCVFLLMARASVCADITAAEGGEAVINCPYGGGYKTSFKYFYKGPYRENTLMLKSDGGESPVFSGRFTLKDDHKARLFTVIIRDLQMNDAGPYYCAA
GWGDSKSIQLNVIRAPKKPKPVQISTSTIHSDQALGSSSHTWPDLSSSAVNPQSTISTFTDHHVSSDSFMIIITVEASVLLLIGLPLLILAVRQRKKTDGLHS
SEFTTSIYQQMTDNPADGDTHTTVCYSCV

>Nilt13(DN2201_c1_g2_i4)
MKFALRVCVFLLMARASVCADITAAEGGEAVINCPYGGGYKTSFKYFYKGPYRENTLMLKSDGGESPVFSGRFTLKDDHKARLFTVIIRDLQMNDAGPYYCAA
GWGDSKSIQLNVIRAPKKPKPVQISTSTIHPNTNSRTDQNSTVTQNESVTMATRTGNTTAEPNEATTRIAGGLGSVLLVLLLCSGTFFILKKRKRKCETALPL
QNVQQNTETDCMYEEILNSDVIIAAEPSSNQTPASDPNTRPENAVYATVTKKRSDSVPGHTHSANRVADTVSDYYGNITSSQQTLNNRTETIYATADLPPKIS
NNEGLIYSVISRK

>Nilt14(ENSDART00000181754)
MNISVLSVVVEAPVTVSGHRGEKLDIRCPYESGYESNSKYFCKGECNFGNKNIIVKSGSPAKDKRFSLTDDKKSRVFTITITDLRTEDEGQYWCAVRRSFTDM
ALTDLYSEIVLMVKQDQENTEGSSVSSFSNTPSYFSSTEVNLQSSSSTDQHNSTGQSHLLQTYSSCAVFMCVCVCVCVCSCSSSV

>Nilt16(ENSDART00000192673)
MMISVFSTVVGAPVTVSGHRGEKLNIRCPYESGYESNSKYFCKGKGILGWKDIIIKSGSPIKDKRFSLSDDKKSRVFTVTITDLRTEDAEQYWCAVERSLPLT
DLYSKVLLKVKQSKTCILIFYNVLFVCFAPHTDLGSVAGGLGSVLLVLLLCSGTFLILKKRKRKCETGNSVSTDCVYENVLDSDVIIAAASSSNQTPASDLNT
HPVITVYSTVTNQQPDLHSGHTQSANQVTDTDCDNYDNITSSEMYVTTTRPQNNTIDDAAIYSLIKRQ

>Nilt17(ENSDART00000193506)
MRDRVCVRKNVWKCTVVEARRTVSGHRGEKLDIRCPYKSGYESYSKYFCKGSCFTGFKDTVVESGSPAIDERFSLTDDKKNRVFTITITDLRSADAGQYWCAV
QRSLLIEDLYSEIVLMVKQGKRSILISLVKLSSTSLIVLNVLFVCFVPHTDLGSVAGGLGSVLLVLLLCSGTFLILKKRKRKCETGNSVSTDCTYENLPKCDV
IIAAKSSSNQMPASDLNTRAANAVYSKVTNQQSELHSGHTHSANQKADSDFDCYANNTNYTTHNAPVSISLLKRKHSGVK

>Nilt18(ENSDART00000187223)
MIISVFSAVVGAPVTASGHRGEKLDIRCSYDPGYESNSKYFCKGKCTPGFKDIIVESGSSPKDKRFSLTDDTTNKVFTVTITDLRTEDAGQYWCAVEKSVFDV
YSEIVLMVKQDQENTEGSSVSSFSNTPSYFSSTEVNLQSSSSTDQHNSTGQSHLLQTYSSCAVFMCVCVCVCVCSCSSSV

>Nilt19(ENSDART00000169976)
MIISVFSAVVGAPVTASGHRGEKLDIRCSYESGYESNSKYFCKGECNFGNKNIIVESGSSAKDKRFSLTDDKKNRVFTITITDLRTEDAGQYWCAVRRGLLIK
DLYSEIVLMVKQDQENTEGSSVSSFSNTPSYFSSTEVNLQSSSSTDQHNSTGQSHLLQTYSSCAVFMCVCVCVCVCSCSSSV

>Nilt21(ENSDART00000152650)
MWDVILLLIYSCLTAVVGAPVTASGHRGEKLDIRCSYESGYESNSKYFCKGKCNFGSKNIIVESGSPAEDDRFSLTDDTINRVFTITITDLRTEDAGQYWCAV
ERSLLIKDLYSEIVLMVKQGTGNTTVQPSTASHTDLGSVAGGLGSVLLVLLLCSGTFLILKKRKRKSLPLQNVQQNTENDCVYENYLDSDVIIAAPSSSNHTP
ASDLNTHPVITVCPTVTNQQPDLHFGHIHSANQVTDTDSYYHRNVASSETIYVTATHPHNNT

>Nilt21(ENSDART00000112472)
MIISVFSAVVGAPVTASGHRGEKLDIRCSYESGYESNSKYFCKGKCNFGSKNIIVESGSPAEDDRFSLTDDTINRVFTITITDLRTEDAGQYWCAVERSLLIK
DLYSEIVLMVKQGKRFILMSLVKLTSHTDLGSVAGGLGSVLLVLLLCSGTFLILKKRKRKCETGNSVSNDCVYENYLDSDVIIAAPSSSNHTPASDLNTHPVI
TVCPTVTNQQPDLHFGHIHSANQVTDTDSYYHRNVASSETIYVTATHPHNNT

>Nilt23(ENSDART00000152507)
FFVVGVLSSISVTGYSGGGVTITCKYAAQYKTKAKYFCKGQWLSTCSDQIKTEIKNKWVQSGRFSLYDNTNAAVFNVTIRNLTEQDSGTYYCGVDLQISDIYT
KVNLKVITVNNTFAKPFSTTTEPKMTLSSEHSTSDLTLSSPSSSNGSSQIVGVWVSVFVILLIIGFGLSIIFFWKRRQTQGSNSASITPATGNSEAVPQMPHF
YEEINNTRPQTDCRTAQLPTISSAACT

>Nilt24(ENSDART00000127091)
MKIIWTFTLLMIPGVLSSISVTGYSGGGVTITCKYDEQYKTNSKYFCKGKYIPTGPDLIKTEIKNKWVKKDKYFLYDNTTAAVFTVTIINLTEKDSGTYYCGV
DIHYHTDIYTEVNLKVITAACCQKSISLSAAAGGSVNISCRYPQSHTADVKFICRRSGADLCAEETRSEEESGGRKAEGKIQLYDDREEQLLTATISHESPEH
SAEYWCGVQSAGHKSFITRVFINFTDAETPPSSPPTSSTSLSKSASSPPNSSSPPPPSSSSASLSDSSSSSAKTSLSSSPKTSPYALIPASPNTSAGSSLIIP
LVLVVLLLIIAALLLLFLYKKHQTKGGDSQTGPGNTAVVSHTACDYEEIKDTHKQLPTNPSESSSAVYSTAELPTKPSESSSAVYATAQLPTNPSESFSSVYA
TAQLPNNPSDSCLYSTVQEASGDSQICISSAEGLNYSVVNFHKTPDLRNSQECSEYAAVNHNFTA
>Nilt24(XM_021478185.1)
MKIIWTFTLLMIPAACCQKSISLSAAAGGSVNISCRYPQSHTADVKFICRRSGADLCAEETRSEEESGGRKAEGKIQLYDDREEQLLTATISHESPEHSAEYW
CGVQSAGHKSFITRVFINFTDAGQTLETIRKVLSSYVCEILSDLIVLFSNAAETPPSSPPTSSTSLSKSASSPPNSSSPPPPSSSSASLSDSSSSSAKTSLSS
SPKTSPYALIPASPNTSAGSSLIIPLVLVVLLLIIAALLLLFLYKKHQTKGGDSQTGPGNTAVVSHTACDYEEIKDTHKQLPTNPSESSSAVYSTAELPTKPS
ESSSAVYATAQLPTNPSESFSSVYATAQLPNNPSDSCLYSTVQEASGDSQICISSAEGLNYSVVNFHKTPDLRNSQECSEYAAVNHNFTA

>Nilt25(XM_009295626.3)
MKIIWTFTLLMIPGVMSSISVTGYSGGGVTITCKYDEGYETYVKYFCKGQWSECTDRIRTGTKDTWVQSGRFSLYDDTTAAVFTVTIRDLTEEDSDMYYCGTD
IYGKVDSATEVNLKVITAACCQKSISLSAAAGGSVNISCRYPQSHTADVKFICRRSGADLCAEETRSEEESGGRKAEGKIQLYDDREEQLLTATISHESPEHS
AEYWCGVQSAGHKSFITGVFIHFTDAESTSPRPSSSPSSPSSSSSSPPSSSSSLVESPLQRMCVRK

>Nilt26(XM_017355936.2)
MEKYRRISDKTGVLSSISVTGYSGVTITCKYDEEYETNEKYFCKGKWSECEDQIRTEMKHHWIRSGRFSLFDDTTAAVFTVTIRNLRKQDSGIYHCGIKRFAF
DHGTKVNLKVITEFGNQVNTNVGTVRRHSGESITIDYTYEEKHKSREKSVCKIGEYLCETLISISRPAEKKDSGRFSIHDDRSAGLLRVFIRELNVQDSGEYR
IKVRHSEDYSFFSEFELVITDDAPKTTSSSPSSSLSSSSSPSSSSYISTLQTPLSTTVSERSIVTSQFNTITDSSLIIPLVLVLLLLIIAALLLLFLYKKHQT
KAGGDSTSQTAHGNTEAVSNIGCDYKEIKDTHKQLPRSPSESSSTVYATAQLPTNPSDFCIYSNVQEASGDSQICISSAEDANYSVVNFHKKSDCPDSISLRN
HQECCEYAAVNHHLTA

>Nilt27(DN4836_c0_g5_i1)
MKIIWTFTLLMIPGVLSSINVTGYSGGGVIITCKYDEQYKTNAKYFCKGQWLSCSELIKTDTKSATKWVQSGRFSLYDNTTAAVFTVTIRDLTRQIYGTYHCG
VEVYGSDPNIEVNLKLVTGSQMGKVRGYSGGNIIIDYRYDKQHKNSKKYVCKTGENHCLSLISIYGGAEWEHSGRFSIRDNRSA

>Nilt28(XM_021478187.1)
MKIIWTFTLLMIPGVVSSMSVIGYSGGGVMITCKYDKGFETYPKYFCKEQWISPCSELIRTEINSEEKWVQSGRFSLIDNKTASGLTVTIRDLTEQDSGTYYC
GVEIYGLDPGTKVNLEVITAGPRIGTVRGYSGGHVIINYSYAIQHKNSQKHVCKTGENLCLSLISISGGAEWEHSGRFSIHDNRSAGLLRVFIRELNVQDSGE
YRIIVRASEDYSFFSEFDLVITDAACCQKSISLSKNVTLVF

>Nilt29(XM_017355937.1)
MKIIWTFTLLMIPGVVSSMSVIGYSGGGVMITCKYDRGFETYPKYFCKEQWISPCSELIRTEINSEEKWVQSRRFSLIDNTTAAFLTVTIRDLTEQDSGTYYC
GVETYGPDLGTKVNLEVITGPRIGTVRGYSGGHVIINSSYEIQHKNSQKHVCKTGENHCLSLININAAAEWKDSGRFSIHDDRSADLLRVFIRELNVQDSGEY
RIIVRESEDYSFFSEFELVVTDGEPLSVIPAFVDVLDVLKIWL

>Nilt29(DN4836_c0_g1_i1)
MKIIWTFSLLMIPGMVSSMSVIGYSGGGVMITCKYDRGFETYPKYFCKEQWISSCSELIRTEINSEEKWVQSRRFSLIDNTTAAFLTVTIRDLTEQDSGTYY

>Nilt30(XM_017357072.2)
MKILLIFILTLQLISGQNSSLSATGYSGGSLMLDSGKLWISDSAKYLAKLPNWRTIINETKHERWINEGRFTLFRNNNGNLMIFIRDLHTDDAGEYWLGALYN
WGADLTLHVEEDSCCASKRVIMTVGETGSFSCDYSHNHINDPKIIFKEEQHSIKMIYSTLNKKDRFNISDDGLRNSFSVSITAVTADDGGVYLCGVWVNKHSY
SYYIINTVHLQIMMSPVVRREGESAQIFCPYDSIYQSKSKSLCKGKCSTRDTRPLDETVREEKERLTLHEDVTASVFTGTITGLTAEDAGKYWCAVTLDRELN
YLYTHLMVIIKQELSLTKYEGDDVSIQCRHHDGDQKSFCRAHEASACVKDGDSLETIRDARFSISDEASAGVFTVNITDLRADDSGVYWCGAHAVTKVHLNIK
QDFSIIIIIIIIIVCVIVLLISGFTLTKRKSRGKRRSDAPLRSIYCTVTHNPL

>Nilt40(GDQH01034272.1)
IINSTWKQVRFSWFNIRDKRHFNVIITAVKADDGGVYLCGVWINEHSYSYYIINTVHLQIITKVGVSTVIGYSGAALMIKCEHPQHKTKTRYICKESSAECSE
EWMKNGDVSVDEDSRAGVLMVFFRELKAADAGTYRCGVKDSEYTERFTQLPVHLGGDVNISCQMAKKMKVYFCKEDNQRCQIMRASKEPQMTPGERVYNVSVR
DAGVYWCGAETRDTHLTFISLTTKIQLSVIMSPVVRREGESAQIFCPYDSIYQSKSKSLCKGKCSTRDTRPLDETVREEKERLTLHEDVTASVFTGTITGLTA
EDAGKYWCAVTLDRELNYLYTHLMVIIKQELSLTKYEGDDVSIQCRHHDGDQKSFCRAHEASACVKDGDSLETIRDARFSISDEASAGVFTVNITDLRADDSG
VYWCGAHLITKVNLTVIKDFSKIVIIITGVCVIVLLICGFTLTVCKLRRKRTEITSTSDKGKRVKNTASSDSRRDPGSAEMTSPDEGSSDELLYASVTFQKHR
ESLSEAPVAFRTITGYSGGSLM

>Nilt40(XM_021475896.1)
VHLGGDVNISCQMAKKMKVYFCKEDNQRCQIMRASKEPQMTPGERVYNVSVRDAGVYWCGAETRDTHLTFISLTTKIQLSVIMSPVVRREGESAQIFCPYDSI
YQSKSKSLCKGKCSTRDTRPLDETVREEKERLTLHEDVTASVFTGTITGLTAEDAGKYWCAVTLDRELNYLYTHLMVIIKQELSLTKYEGDDVSIQCRHHDGD
QKSFCRAHEASACVKDGDSLETIRDARFSISDEASAGVFTVNITDLRADDSGVYWCGAHLITKVNLTVIKGKIRAHYS

>Nilt48
SKRVTVNSGETAVFSCEYSHNQIHHAKVIFKEERDSVKSVIHTTARIREEGRVHMSDDRQRNVLSVSITAVTAGDGGVYLCGVWVNKHSYSYYITNTVHLQIM
DVSMIFIIIVCVSVFFIGAFTLAVCKLGQEKQSDAAVHTPLPTTPSDETIVIFQKHAETLRVAADAFMTDEVYSDYTMVIYYTELN
>Nilt51(XM_021478188.1)
MKILLIFILTLQLISGQISSLSATGYSGGSLMLDSGKLWVSDSAKYLAKLPDWRTIINDTKHERWINEGRFTLFKNNDGNLMIFIRDLNTDDAGRYAVDVEQG
KRLYMILNVEEDSCCNMSKRVTVNSGETAVFSCEYSHNQINHAKVILKPERDSIEMIYSTLNKQDRFSISDDRQRNSFSVSITAVTAGDGGVYLCGVWVNKHS
YSYYITNTVHLQIMGQEKQSDAAVHTPLPTTPSDETIVIFQKHAETLRVAADAFMTDEVYSDYTMVIYYTELN

>Nilt52(DN78550_c0_g1_i1)
MKILLIFFVFTLQLMSAEVRYIDVTGYLGGNLLVSSWEPWHESDAKFMEKLRGGIIINNRKQRQWINEGRFTLFYTDYRNLVIYIKELVTEDAGTYRIKVKGK
WSINMILSVKKDSCCEVPKTVMVNNG

>Nilt55(XM_017356846.2)
MCVCLRLVFIVHCFLNVLLLGQEALNRGSVFLLTGHMGDEAVSVTGVAGGSVIFNFQYKDKISVERFCRDADDECIQAQSHLWLPEEKLTMTDNKSVNYLSVL
IRNLTVNDSGTYEWKDNKDWSKEVKLVVQQEPCCGRSEEQTAYVQGTAVIHCRYPKTNHDYSKQLFKEQNGSLEAIFTSREASDPRFSINLNHGEEHFFTVTL
TNVSRHDEGLYFCAAHGKMSIRYSLLFTEIQLHVTDAPVQSDLSVLIITVSVCVILLFAAGLSLFVLRRRCTKTKGSKSSDQRSTVEIDEINSSVYYSSIKEQ
CVQKPVSMNTVYATAQLPTVLSDSDPYTLAMKS

>Nilt55(XM_009295658.3)
MMMTMMMMMMMKTFLFFTIFLIEGHMGDEAVSVTGVAGGSVIFNFQYKDKISVERFCRDADDECIQAQSHLWLPEEKLTMTDNKSVNYLSVLIRNLTVNDSG
TYEWKDNKDWSKEVKLVVQQEPCCGRSEEQTAYVQGTAVIHCRYPKTNHDYSKQLFKEQNGSLEAIFTSREASDPRFSINLNHGEEHFFTVTLTNVSRHDEGL
YFCAAHGKMSIRYSLLFTEIQLHVTDAPVQSDLSVLIITVSVCVILLFAAGLSLFVLRRRCTKTKGSKSSDQRSTVEIDEINSSVYYSSIKEQCVQKPVSMNT
VYATAQLPTVLSDSDPYTLAMKS

>Nilt56(XM_003197773.5,ENSDART00000128873)
MIHISEDKLLIFTLLILMTVVYSEMKTFTGHEGGKVEIRCPYNAKYEEMEKYLCRGECPIYDKDIPVESGSEPVDKRFSLTDNRTAHIFTITITDLRTDDQGK
YWCAVKTGIGSFDKYTEIYLEIKQVSRVSGEPGKHLSITCSYTRDLKELIRFLCKGSDPSDCKNIIKVSSKTKTNGRFSLTDDSETVFTVNIRDLTEEDSGIY
WCGAAEKQAQEYTWISAIDLHITEPTSAPRLPKNTSTASFHTSKPADPASSRPVTPSSSASSSSISMFLNASTNAASSHTPLGFTAFIMLMVVVMLTVFGLSL
FLYLKRKQKKKGAQTKDDIYYQPENLPNGNTETTREASHGVSDYEEISSSLNNPIYSSVSPAFSKQDVSVYAVAQLPSSPSDHLNYSTVRFSASHLSDRTSGG
LDTCDYATVNV

>Nilt57(DN34828_c0_g1_i1)
MRSAFTIFITLFLISGQGLCFKVHGCTGGSLLFKCMYSNQKSSDVENDGAYFSRDNPYKMVINTKIQSCWSHNGRFAVYDDEIRKFILVLIKNLSREDDGTYT
CGHDHKWNHNVELVVESESSCGTSVIQAAYAGQETTIICKYQDKFKKNTKVFYTLNGDPVHMLNSSSQSSVKRFSLSDSHKDHFNVTISSVSAEDDGDYLCGV
ERSDDQRSSSITHNTFIQKIHLNVTNKITSVQVQAYIGRSVFIKCKFLQKFKENKKFFMRAESQKFMLDEQNQWIHHDNVHMYDNTTEGHLKVFISDLSAANE
GTYRCGVNIDRDDLYTEVKLNVNKAGDFHSSSEAAFIGESVKLTCIHREKNTVNHICKENRDKICQNIVSSNKGRVQISDSTDGVFTVSISNVSVRDAGVYWC
GAKTRDTHLTFISLTTKLELTLTMQPVIGREGDSAEIRCPYEDHNLSEIKYFYKGKCFTQEAPIIRSDEGHIKKPNISMKDNTELNVFTVNISSLRAEDAGIY
CCVVRDMFIISTELMIIMKEVPVIHKASVRGSASISCKCIRKHTQHQRFFCKGNQPNICVHAGVLVSFNKPANSRFSQTDECSAGVFTVNISQLTEEDSGKYW
CGEESSGSFLFTEVHLNVNTETNMPDPNSANSGLSVIVVVSVCLILIAVCLLLLFFKKKCYSKQVRTGITSSADRRQTADHETAQEEIQDSSPLYSVQLPTIP
SDGLLYASVSFQKHGESLGEAAVAFNKDETHCDYASVNHRLTHN

>Nilt58(ENSDART00000123831)
VFRSAVSVVNSISVTGYSGGGVTITCKYDRQYKKNTKYFCKGQKFLSCSKLIKTEAKSDNKWVQKDRYSLFHNTTAADFTVTIRNLTVWDSGTYYCGVEVYGL
DPSTKVNLIIIAVDDRFTKPFSITTEPEFTLNSEHSTSDLISSAPSSSNGLSQIIGVSVTVILLIFGFVSSIVFCKRRQRKGSNSASITPATGNSEAVPQMPH
FYEEINHTRPQTDCRTAQLPTISSAACSTVYATQQLPTNPSSCSTVYATPHQPANPSAPSSIVYSTPQFPTNPSAFCSTVYATQQLPTNPSASSSTVYATPHQ
PTNPSASSNVVHTAD

>Nilt58(ENSDART00000156066)
VFRSAVSVVNSISVTGYSGGGVTITCKYDRQYKKNTKYFCKGQKFLSCSKLIKTEAKSDNKWVQKDRYSLFHNTTAADFTVTIRNLTVWDSGTYYCGVEVYGL
DPSTKVNLIIIAGLSQIIGVSVTVILLIFGFVSSIVFCKRRQRKGSSLIIGGSVTVILLISGFVSSIILFCKRRQTRGSNSASITPATGNSEAVPQMPHFYEE
INHTRPQTDCRTAQLPTISSAACSTVYATQQLPTNPSSCSTVYATPHQPANPSAPSSIVYSTPQFPTNPSAFCSTVYATQQLPTNPSASSSTVYATPHQPTNP
SASSNVVHTAD

>Nilt61(XM_005169735.1)
MKIIWTFTLLMIPGVLSSISVTGYSKGGVTITCKYDKGYETYVKYFCKGEWSGCTDQIKTATQDKWIKSGRFSLYDNTKEAHLIVTIKDLREQDSDIYYCGTE
VYGYDLSTGVNLKVIADSFSNGGAKCFRHNFIFTIFMFTIFIFTILIFITIIFISGSISTAENVC

>Nilt63(XM_005169736.1)
MKIIWTFTLLMIPGVLSSISVTGYSGGGVTIRCKYDEEYETNAKYFCRGEWPECTDQIRTEMKHQWIRSGRFSLFDDTRAAVFTVTIRNLRKQDSGMYYCGIK
IFGPDHDTKVNLKVITEFRNQVNTNVGTVRRHSGGNITIDYTYEKQHKSREKSVCKIGEYQCETLISISRPAEKDSGRFSIHDDRSAGLLHVFIRELNVQDSG
EYRIIVRHSEDYSFFSEFDLVITDDASKTTSSSLSSSSSPSSSSYISTLQTPLSTTVSEKSIITSQFNTITDSSLIIPLVLVLLLLIIAALLLLFLYKKHQTK
GGDSTSQTAHGNTEAVSNIGCDYKEIKDTHKQLPRSPSESSSTVYATAQLPTNPSDFCIYSNVQEASGDSQICISSAEDAHYSVVNFHKKPDCPDSISLRNHQ
ECCEYAAVNHHLTA

>Nilt66(XM_005169737.1)
MAKLGMSIVSSFAGAALMIKCEHPQHQTKTRYICKESSAGCSGEWRKNGDVSVDEDTRAGVLMVFFRELKAADAGTYRCGVKDSEYTERFTQLQLKVRHDGKY
QEVMNESVQLGGEVKISCQIPDEQKVHFCKEDDHKSCQIMRASEDPQLTPAGSVGNKERVFTVSISNVSVRDAGVYWCGAETRDTHLTFISLTTKIQLSVIMS
PVVRREGESAQIFCPYDSIYQSKSKSLCKGKCSTRDTRPLDETVREEKERLTLHEDVTASVFTGTITGLTAEDAGKYWCAVTLDRELNYLYTHLMVIIKQELS
LTKYEGDDVSIQCRHHDGDQKSFCRAHEASACVKDGDSLETIRDARFSISDEASAGVFTVNITDLRADDSGVYWCGAHLITKVNLTVKIDLIDVLRKNFTIIT
IILVFITAENVCQEIIHIKGV

>Nilt67(XM_005169733.1)
MKIIWTFTLLMIPGMVSSMSVIGYSGGGVMITCKYDRGFETYPKYFCKEQWISPCSELIRTEINSEEKWVQSGRFSLIDNKTAAGLTVTIRDLTEQDSGTYYC
GVETYGPDLGTKVNLEVITGPRIGTVRGYSGGHVIINYSYAIQHKNSQKHVCKTGENHCLSLISISGAAEWKHSGRFSIHDDRSAGLLRVFIRELNVQDSGEY
RIIVRASEDYSFFSEFELVVTDDASETTSSSSPPPPPSSSSSLVESPLQRMCVRL

>Nilt68(ENSDART00000121467)
MNCIWILCPSMVSSMSVIGYSGGGVMITCKYDRGFETYPKYFCKEQWISPCSELIRTEINSEEKWVQSGRFSLIDNKTAAGLTVTIRDLTEQDSGTYYCGVET
YGPDLGTKCCYRCWFVVGVVSSMSVIGYSGGGVMITCKYDEQYKTSAKYFCKEQWISPCSELIRINSEEKWVQSGRFSLFDNRTAAGLTVTIRNLTEQDSGTY
HCGVETYGPDPGTKVNLEVITGKNLSSSNISTLQTPLSTTVSENSTLTSQFNTITESSLIIPLVLVLLLLIIAALLLLFLYKKHQTKAGGDSTSQTAHGNTEA
VSNIDCDYEEIKDTHKQLPRSPSDSSSTVYATAQLPTNPSDSYMNYSVVNFHKTPDCPDSVSL

>Nilt68(ENSDART00000124611)
QICLVDLSLLLLFACFNLYVSIICMVSSMSVIGYSGGGVMITCKYDRGFETYPKYFCKEQWISPCSELIRTEINSEEKWVQSGRFSLIDNKTAAGLTVTIRDL
TEQDSGTYYCGVETYGPDLGTKILHCSFNHRVVSSMSVIGYSGGGVMITCKYDEQYKTSAKYFCKEQWISPCSELIRINSEEKWVQSGRFSLFDNRTAAGLTV
TIRNLTEQDSGTYHCGVETYGPDPGTKVNLEVITGKVISSAPVCLTKQGMIRSLNTSTLQTPLSTTVSENSTLTSQFNTITGTHVVLLLLIIAALLLLFLYKK
HQTKAGGDSTSQTAHGNTEAVSNIDCDYEEIKDTHKQLPRSPSDSSSTVYATAQLPTNPSDSYMNYSVVNFHKTPDCPDSVSL

>Nilt69(XM_005169738.1)
MTVPFPCSSSAVSPVVRREGESAQIFCPYDSIYQSKSKSLCKGKCSTRDTRPLDETVREEKERLTLHEDVTASVFTGTITGLTAEDAGKYWCAVTLDRELNYL
YTHLMVIIKQELSLTKYEGDDVSIQCRHHDGDQKSFCRAHEASACVKDGDSLETIRDARFSISDEASAGVFTVNITDLRADDTGVYWCGAHAVTKVHLNIKQD
FSRIIIIIIIVVCVIVLLISGFTLTKRKSRGKRRSDAPLRSIYCTVTHNPL

>Nilt70(XM_005169734.1)
MLDSGKRWFSDSAKYLAKLPEWRTIINDRKHERWINEGRFTLFKNTDANLMIFIRDLHTDDAGRYAVDVEQGNRIYIILNVGEDSCCNVSKRVTVNSGETAVF
SCEYSHNQIHHAKVIFKEERDSVKSVIHTTARIREEGRVHMSDDRQRNVLSVSITAVTADDGGVYLCGVWVRENSYNYSTLHTINLHVITKVGVSTVIGYSGA
ALMIKCEHPQQKTNTKYICKESSAGCSEEWRKNGDVSVDEDTRAGVLMVFFRELKAADAGTYRCGVKDSEYTERFTQLQLIIRHDGEYPKVMNESVHLGGEVK
ISCQIPEEQNKNYFCKEDDHKSCQIRRASEDPQLTPAGSVGSEERVFTVSISNVSVRDAGVYWCGAETRDTHLTFISLTTKIQLSVISDDFSMKPEISIIILT
GVCVIALLAVGFPLTVCIIGHKQTGKTCTAEKKTREKTTSSDSGHDQTVSPDGASSAGLLYASVTFQKHQESLSEAGVTFRTSQVYSDYTTVVLPQ

>Nilt70(ENSDART00000125051)
MLDSGKRWFSDSAKYLAKLPEWRTIINDRKHERWINEGRFTLFKNTDANLMIFIRDLHTDDAGRYAVDVEQGNRIYIILNVGEDSCCNVSKRVTVNSGETAVF
SCEYSHNQIHHAKVIFKEERDSVKSVIHTTARIREEGRVHMSDDRQRNVLSVSITAVTADDGGVYLCGVWVRENSYNYSTLHTINLHVITKVGVSTVIGYSGA
ALMIKCEHPQQKTNTKYICKESSAGCSEEWRKNGDVSVDEDTRAGVLMVFFRELKAADAGTYRCGVKDSEYTERFTQLQLIIRHDGEYPKVMNESVHLGGEVK
ISCQIPEEQNKNYFCKEDDHKSCQIRRASEDPQLTPAGSVGSEERVFTVSISNVSVRDAGVYWCGAETRDTHLTFISLTTKIQLSVISDG

>Nilt75(XM_005169739.1)
MIKCKHPQHKTKTRYICKESSAGCSEEWRKNGDVSVDDDSRAGVLMVFFRELKAADAGTYRCGVKDSEYTERFTQLQLNIRHDAKYPKVMKESVQLDGEVKIS
CQILEEQKVHFSYFCKEDDHKKCQKTRASEEPQRTPAGSVGSEERVFTVSISNVSVRDAGVYWCGAETRDTHLTFISLTTKIQLSIISESIQMKTKVGVSTVI
GSSGATLMIKCEHPQHKTNTKYICKESSAGCSEEWRKNGDVSVDDDSRAGVLMVFFRELKAADAGTYRCGVKDSEYTERFTQLQLKVRHDAIFINKTTRLGGE
VNISCQTPEEHIAYFCKEDDNDCYNRIRQNFSSAEEPQMNPAEFVSISNVSVRDAGVYWCGAETRDTHLTFISLTTKIQLSVIMSPVVRREGESAQIFCPYNS
IYQSKSKSLCKGKCSTRDTRPLDETVREEKERLTLHEDVTASVFTGTITGLTAEDAGKYWCAVTLDRELNYLYTHLMVIIKQELSLTKYEGDDVSIQCRHHDG
DQKSFCRAHEASACVKDGDSLETIRDARFSISDEASAGVFTVNITDLRADDSGVYWCGAHVITKVNLTVKKSVFARTLISCVSVTLLLTGASVLLFYISGFIK
KRDRHSSSVTETSRGEASQAACKYENITYVKQHLASGTEETVLYSTVNRPTDSSDAHEMLYSTVQLPKNSSDEILYSSVHFQKHKESDSEAEVFLNDELYCDY
TTVRQQS

>Nilt78(XM_003197772.2)
MKTFTGHEGGKVEIQCPYSESAYEGNQKYLCRGECPIKPFLFQNIDIPVESGSAPVDKRFSLTDNRTAHIFTITITDLRTADQGKYWCAVKTGLVKFDDYKQI
YLEIKQVSRVSGEPGKHLSITCSYTRDLKELIRFLCKGSNPSHCKNIIKVSSETKTNGRFSLTDDSERNFTVNISDLTEEDSGIYWCGAEDNQTQEYTWISAV
DLHITDATSERRLPKNTSTASFHTSKPADTASSRPVTPSSPASSSSSTSMLSSWKRLLCTVFIILITSVMLTVFGLSVYIFLRWKPQTKGADRRNAAHDLTEK
LLNTEEAPGDIYTVCHYEELTHTDDHPAGLGSGRPVFGEQDPLTCSTIIFNTGLHSDMTSD

>Nilt79(XM_003197773.2)
MIHISEDKLLIFTLLILMTVVDSEMNTFTGHEGRKVKIKCPYDAKFEKMEKYLCRGKCSIGYKDIPVESGSEPKDKRFSLTDNRTAHIFTITITDLRTADQGK
YWCAVKTRLGKFDDYKEIYLEIKQVSRVSGEPGKHLNITCSYTRDLKELIRFLCKGSDPSDCKKMIKVSSETNTTGRFSLTDDSERNFTVNIRDLTEEDSGIY
WCGAEEKQTRKHTWISATDLQISEATSERRLPKNTTAASFHTSKPEDPASSRPVTPSSSASSSSISMFLNASTNAASSNTPLGFTAFIMLMVVVMLTVFGLSL
FLYLKRKQKKKGAQIKDDVYYQPENLPNGNTETTREATHGVSDYEEISSSLNNPIYSSVSPAFSKQDASVYAVAQLPSSPSDHLNYSTVRFSASHHSDRTSGG
LDTCDYATVNV

>CG1_transcript_2(DN4836_c0_g4_i1)
MKIIWTFTLLMIPGVLSSISVTGYSGGGVTIRCKYDEEYETNAKYFCRGEWPECTDQIRTEMKHHWIRSGRFSLFDDTRAAVFTVTIRNLRKQDSGMYYCGIK
IFGPDHDTKVNLKVITEFGNQVNTNVGTMRRHSGGNITIEYTYEEKHKSREKSVCKIGEYQCETLISISRPAKKKDGGRFSIYDNRSAGLLRVFIRELNVQDS
GEYRITVRVSEDYSFFSEFELDITDDASKTTSSSLSSSS

#### Figure S13. Sequence diversity of zebrafish NILT proteins.

Predicted NILT proteins encoded by transcripts listed in **Supplemental Table S3** and graphically displayed in **Figure 6** are shown with sequence identifiers in parentheses. Ig domains with the $Cx_3C$ motif are shaded blue. Ig domains with the $Cx_6C$ motif are shaded red. Ig domains with the $Cx_7C$ motif are shaded orange. Ig domains defined as atypical are shaded purple. ITIM and ITIM-like (itim) sequences are shaded black and gray, respectively. The ITAM sequence in Nilt10 is underlined. TVYAT sequences in Nilt1, Nilt26, Nilt55, Nilt58, Nilt63 and Nilt68 are double underlined. Leader sequences and transmembrane domains are indicated with boxed text. Lower case letters indicate protein sequence predicted from genomic sequence. Transcripts encoding only a single Ig domain are excluded from this list, but can be found in **Supplemental Table S3**.

```
Dare-Nilt10     DQMYTELNT--RQTDLYHSLTTD
Cyca-NILT1      DQIYTELNA-SRQSDVYQSLRTD
Onmy-NILT1      ESVYQDITP--Y-DNIYQSLNIT
Onmy-NILT2      DGDYEEINEQPLHSNIYANLPTS

Dare-Dap12      ESPYQELYG--VQSDIYSDLQQY
Mumu-Dap12      ESPYQELQG--QRPEVYSDLNTQ

Dare-CD3z.1     ESHYQAIDK--LGKSEYATANRT
Dare-CD3z.2     DETYTPLTR--KGDDTYRELETK
Dare-CD3z.3     DQLYQGLSS--VTKDTYDSLQMQ
Dare-CD3zl.1    EPVYTDLDL-PQVGSDYQQLDRP
Dare-CD3zl.2    ETHYQELRA--HASDEYQQIGTK
Mumu-CD3z.1     NQLYNELNL-GRR-EEYDVLEKK
Mumu-CD3z.2     EGVYNALQK-DKMAEAYSEIGTK

Dare-FcRg       EGVYEGLKP--HETDTYETIKMK
Dare-FcRgl      GDLYQDLGR--RDADTYDTLHGM
Mumu-Fcrg       DAVYTGLNT--RSQETYETLKHE
```

#### Figure S14. ITAM sequences.

ITAM sequences from zebrafish (Dare) Nilt10, carp (Cyca) NILT1 (CAH19212.1), trout (Onmy) NILT1 and NILT2 (XP_021475862.2 and CAQ77254.1), were aligned with ITAM sequences from Dap12 from zebrafish and mouse (Mumu) (ABO61032.1 and NP_035792), CD3ζ (CD3z) and CD3ζ-like (CD3zl) sequences from zebrafish and mouse (ABS18393.1, ABO61034.1 and NP_112439), and FcRγ (FcRg) and FcRγ-like (FcRgl) sequences from zebrafish and mouse (ABO61033.1, ABS18392.1 and NP_034315). If a protein possessed multiple ITAMs, they are indicated numerically after the protein symbol (e.g. the second ITAM in zebrafish CD3ζ is listed as Dare-CD3z.2).

#### ITIMs and itims

| | |
|---|---|
| Nilt1.1 | DLYGDV |
| Nilt1.2 | TVYATV |
| Nilt2 | CHYEEL |
| Nilt3.1 | RAYACV |
| Nilt3.2 | PIYQNI |
| Nilt3.3 | AIYGNV |
| Nilt4.1 | SIYENL |
| Nilt4.2 | EDYGNL |
| Nilt5.1 | GDYEEI |
| Nilt5.2 | STYCNV |
| Nilt5.3 | PPYSLL |
| Nilt9.1 | LNYSSV |
| Nilt9.2 | THYTSV |
| Nilt9.3 | VIYSTV |
| Nilt13.1 | CMYEEI |
| Nilt13.2 | AVYATV |
| Nilt13.3 | DYYGNI |
| Nilt13.4 | LIYSVI |
| Nilt17.1 | CTYENL |
| Nilt17.2 | AVYSKV |
| Nilt23 | HFYEEI |
| Nilt24.1 | CDYEEI |
| Nilt24.2 | CLYSTV |
| Nilt24.3 | LNYSVV |
| Nilt24.4 | SEYAAV |
| Nilt26.1 | CDYKEI |
| Nilt26.2 | CIYSNV |
| Nilt26.3 | ANYSVV |
| Nilt26.4 | CEYAAV |
| Nilt30 | SIYCTV |
| Nilt40 | LLYASV |
| Nilt48.1 | SDYTMV |
| Nilt48.2 | IYYTEL |
| Nilt55 | VYYSSI |
| Nilt56.1 | SDYEEI |
| Nilt56.2 | PIYSSV |
| Nilt56.3 | LNYSTV |
| Nilt56.4 | CDYATV |
| Nilt57.1 | LLYASV |
| Nilt57.2 | CDYASV |
| Nilt58 | HFYEEI |
| Nilt63.1 | CDYKEI |
| Nilt63.2 | CIYSNV |
| Nilt63.3 | AHYSVV |
| Nilt63.4 | CEYAAV |
| Nilt68.1 | CDYEEI |
| Nilt68.2 | MNYSVV |
| Nilt69 | SIYCTV |
| Nilt70.1 | LLYASV |
| Nilt70.2 | SDYTTV |
| Nilt75.1 | CKYENI |
| Nilt75.2 | VLYSTV |
| Nilt75.3 | MLYSTV |
| Nilt75.4 | ILYSSV |
| Nilt75.5 | CDYTTV |
| Nilt78 | CHYEEL |
| Nilt79.1 | SDYEEI |
| Nilt79.2 | PIYSSV |
| Nilt79.3 | LNYSTV |
| Nilt79.4 | CDYATV |

#### ITIMs only

| | |
|---|---|
| Nilt4.1 | SIYENL |
| Nilt5.2 | STYCNV |
| Nilt9.1 | LNYSSV |
| Nilt9.3 | VIYSTV |
| Nilt13.4 | LIYSVI |
| Nilt24.3 | LNYSVV |
| Nilt24.4 | SEYAAV |
| Nilt30 | SIYCTV |
| Nilt40 | LLYASV |
| Nilt48.1 | SDYTMV |
| Nilt48.2 | IYYTEL |
| Nilt55 | VYYSSI |
| Nilt56.1 | SDYEEI |
| Nilt56.3 | LNYSTV |
| Nilt57.1 | LLYASV |
| Nilt69 | SIYCTV |
| Nilt70.1 | LLYASV |
| Nilt70.2 | SDYTTV |
| Nilt75.2 | VLYSTV |
| Nilt75.4 | ILYSSV |
| Nilt79.1 | SDYEEI |
| Nilt79.3 | LNYSTV |

#### itims only

| | |
|---|---|
| Nilt1.1 | DLYGDV |
| Nilt1.2 | TVYATV |
| Nilt2 | CHYEEL |
| Nilt3.1 | RAYACV |
| Nilt3.2 | PIYQNI |
| Nilt3.3 | AIYGNV |
| Nilt4.2 | EDYGNL |
| Nilt5.1 | GDYEEI |
| Nilt5.3 | PPYSLL |
| Nilt9.2 | THYTSV |
| Nilt13.1 | CMYEEI |
| Nilt13.2 | AVYATV |
| Nilt13.3 | DYYGNI |
| Nilt17.1 | CTYENL |
| Nilt17.2 | AVYSKV |
| Nilt23 | HFYEEI |
| Nilt24.1 | CDYEEI |
| Nilt24.2 | CLYSTV |
| Nilt26.1 | CDYKEI |
| Nilt26.2 | CIYSNV |
| Nilt26.3 | ANYSVV |
| Nilt26.4 | CEYAAV |
| Nilt56.2 | PIYSSV |
| Nilt56.4 | CDYATV |
| Nilt57.2 | CDYASV |
| Nilt58 | HFYEEI |
| Nilt63.1 | CDYKEI |
| Nilt63.2 | CIYSNV |
| Nilt63.3 | AHYSVV |
| Nilt63.4 | CEYAAV |
| Nilt68.1 | CDYEEI |
| Nilt68.2 | MNYSVV |
| Nilt75.1 | CKYENI |
| Nilt75.3 | MLYSTV |
| Nilt75.5 | CDYTTV |
| Nilt78 | CHYEEL |
| Nilt79.2 | PIYSSV |
| Nilt79.4 | CDYATV |

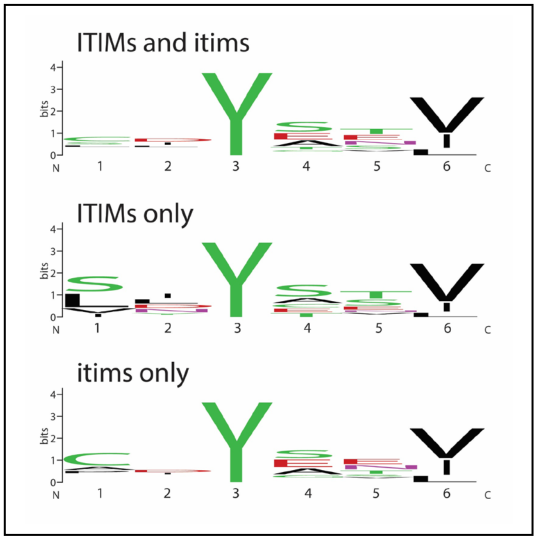

#### Figure S15. ITIM and itim sequences.

ITIM and itim sequences from zebrafish NILTs (listed in **Figure S13**) were aligned jointly (left) or singly as ITIMs (middle) or itims (right). If a NILT possessed multiple ITIMs, they are indicated numerically after the protein symbol (e.g. the second itim in Nilt1 is listed as Nilt1.2). Logo consensus sequences (in box) were generated (weblogo.berkeley.edu) from aligned ITIMs and itims (Crooks et al., 2004). Note that the major difference between NILT ITIMs and itims is in position 1 with ITIMs possessing the consensus S/L/V/I and itims commonly possessing C/A/P.

```
Dare-Nilt1        ASDLHTHSVITVYATVTN*
Dare-Nilt26       LPRSPSESSSTVYATAQLPTNPSDF
Dare-Nilt55       QCVQKPVSMNTVYATAQLPTVLSDS
Dare-Nilt58.1     LPTISSAACSTVYATQQLPTNPSSC
Dare-Nilt58.2     LPTNPS-SCSTVYATPHQPANPSAP
Dare-Nilt58.3     FPTNPSAFCSTVYATQQLPTNPSAS
Dare-Nilt58.4     LPTNPSASSSTVYATPHQPTNPSAS
Dare-Nilt63       LPRSPSESSSTVYATAQLPTNPSDF
Dare-Nilt68       LPRSPSDSSSTVYATAQLPTNPSDS
Sasa-NILT3        LQSSSGGETTTVYATANLPTRPFDS
Sasa-NILT5        LQSSSGGETTTVYATANLPTSPSDS
```

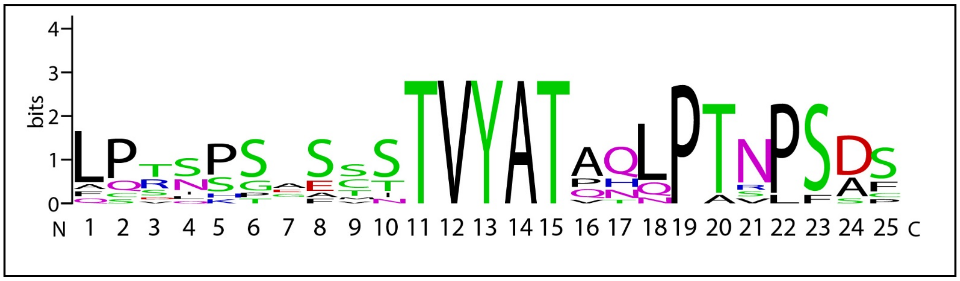

#### Figure S16. TVYAT sequences.

TVYAT peptide sequence motifs from zebrafish (Dare) and Atlantic salmon (Sasa) NILTs were identified by manually scanning all known NILT cytoplasmic tails for similar sequences. In order to identify additional conserved sequences, TVYAT motifs with 10 adjacent residues were aligned (top). Note that the TVYAT motif in Dare-Nilt1 is included within an itim and is at the carboxyl-terminus of the cytoplasmic tail (* = translational stop) and that Dare-Nilt58 possesses 4 TVYAT motifs (numbered 1-4). A logo consensus sequence (bottom, in box) was generated (weblogo.berkeley.edu) from aligned motifs (Crooks et al 2004). Note that with the TVYAT motif in positions 11-15, a proline-threonine dipeptide sequence (PT) in positions 19-20 and a proline-serine dipeptide sequence (PS) in positions 22-23 are highly conserved. The function of this sequence motif, if any, is unknown.

### References.

Crooks GE, Hon G, Chandonia JM, Brenner SE. 2004. WebLogo: A sequence logo generator. *Genome Res*. 14:1188-1190. PMID: 15173120

Dornburg A, Lippi C, Federman S, Moore JA, Warren DL, Iglesias TL, Brandley MC, Watkins-Colwell GJ, Lamb AD, and Jones A. 2016. Disentangling the Influence of Urbanization and Invasion on Endemic Geckos in Tropical Biodiversity Hot Spots: A Case Study of Phyllodactylus Martini (Squamata: Phyllodactylidae) along an Urban Gradient in Curaçao. *Bull Peabody Mus Nat Hist.* 57(2): 147–164. https://doi.org/10.3374/014.057.0209

Hoang DT, Chernomor O, Von Haeseler A, Minh BQ and Vinh LS, 2018. UFBoot2: improving the ultrafast bootstrap approximation. *Mol Biol Evol. 35*(2): 518-522. PMID: 29077904

Kock H, Fischer U. 2008. A novel immunoglobulin-like transcript from rainbow trout with two Ig-like domains and two isoforms. *Mol Immunol*. 45(6):1612-22. PMID: 18035417

Minh BQ, Schmidt HA, Chernomor O, Schrempf D, Woodhams MD, Von Haeseler A, Lanfear R. 2020. IQ-TREE 2: New models and efficient methods for phylogenetic inference in the genomic era. *Mol Biol Evol*. 37(5):1530-4. PMID: 32011700

Østergaard AE, Martin SA, Wang T, Stet RJ, Secombes CJ. 2009. Rainbow trout (*Oncorhynchus mykiss*) possess multiple novel immunoglobulin-like transcripts containing either an ITAM or ITIMs. *Dev Comp Immunol*. 33(4):525-32. PMID: 19013192

Robinson JT, Thorvaldsdóttir H, Winckler W, Guttman M, Lander ES, Getz G, Mesirov JP. 2011. Integrative genomics viewer. *Nat Biotechnol*. 29(1):24-6. PMID: 21221095

Schwartz S, Zhang Z, Frazer KA, Smit A, Riemer C, Bouck J, Gibbs R, Hardison R, Miller W. 2000. PipMaker—a web server for aligning two genomic DNA sequences. *Genome*, 10(4): 577–586. PMID: 10779500

Stet RJ, Hermsen T, Westphal AH, Jukes J, Engelsma M, Lidy Verburg-van Kemenade BM, Dortmans J, Aveiro J, Savelkoul HF. 2005. Novel immunoglobulin-like transcripts in teleost fish encode polymorphic receptors with cytoplasmic ITAM or ITIM and a new structural Ig domain similar to the natural cytotoxicity receptor NKp44. *Immunogenetics*. 57(1-2):77-89. PMID: 15702329

Sievers F, Higgins DG. 2014. Clustal Omega, Accurate Alignment of Very Large Numbers of Sequences. *Methods Mol Biol* 1079: 105–16. PMID: 24170397
